# 5-Deoxyadenosine Salvage by Promiscuous Enzyme Activity leads to Bioactive Deoxy-Sugar Synthesis in *Synechococcus elongatus*

**DOI:** 10.1101/2020.12.30.424818

**Authors:** Johanna Rapp, Pascal Rath, Joachim Kilian, Klaus Brilisauer, Stephanie Grond, Karl Forchhammer

## Abstract

7-Deoxysedoheptulose is an unusual deoxy-sugar, which acts as antimetabolite of the shikimate pathway thereby exhibiting antimicrobial and herbicidal activity. It is produced by the unicellular cyanobacterium *Synechococcus elongatus* PCC 7942, which has a small, stream-lined genome, assumed to be free from gene clusters for secondary metabolite synthesis. In this study, we identified the pathway for the synthesis of 7-deoxysedoheptulose. It originates from 5-deoxyadenosine, a toxic byproduct of radical *S*-adenosylmethionine (SAM) enzymes, present in all domains of life. Thereby we identified a novel 5-deoxyadenosine salvage pathway, which first leads to the synthesis and excretion of 5-deoxyribose and subsequently of 7-deoxysedoheptulose. Remarkably, all reaction steps are conducted by promiscuous enzymes. This is a unique example for the synthesis of a bioactive compound without involving a specific gene cluster. This challenges the view on bioactive molecule synthesis by extending the range of possible compounds beyond the options predicted from secondary metabolite gene clusters.

## Introduction

*S*-Adenosyl-l-methionine (SAM or AdoMet), which is formed by ATP and the amino acid methionine, is an essential cofactor of various enzymatic reactions in all domains of life. SAM can serve as a methyl group donor for the methylation of DNA, RNA and proteins in reactions that release *S*-adenosylhomocysteine (SAH) as byproduct (***Fontecave et al., 2004***). SAM can also serve as an aminopropyldonor for polyamine synthesis, and as a homoserine lactone donor for the synthesis of quorum sensing compound *N*-acetylhomoserine lactone which both result in the release of 5-methylthioadenosine (MTA). Furthermore, SAM is a source of the 5-deoxyadenosylradical (5dAdo^•^), which is formed by the activity of radical SAM enzymes (***Booker and Grove, 2010; Broderick et al., 2014; Fontecave et al., 2004; Sofia et al., 2001; Wang and Frey, 2007***). 5dAdo^•^ is formed by the reductive cleavage of SAM and can abstract a hydrogen atom from its substrate to form a substrate radical as well as 5-deoxyadenosine (5dAdo), which is released as a byproduct (***Marsh et al., 2010; Wang and Frey, 2007***). Radical SAM enzymes, a superfamily with over 100.000 members, are present in all domains of life (***Holliday et al., 2018; Sofia et al., 2001***). They are catalysing various complex chemical reactions, including sulphur insertion, anaerobic oxidations, unusual methylations and ring formations (***Parveen and Cornell, 2011***). Prominent members of this family are, for example, involved in biotin, thiamine and lipoate biosynthesis. Other members are involved in DNA repair or in the biosynthesis of secondary metabolites e.g. antibiotics (***Wang and Frey, 2007***). MTA, SAH and 5dAdo are product inhibitors of these reactions (***Challand et al., 2009; Choi-Rhee and Cronan, 2005; Farrar et al., 2010; Palmer and Downs, 2013; Parveen and Cornell, 2011***). Therefore, and because of the high bioenergetic costs of these compounds, salvage pathways are necessary. SAH is rescued via the methionine cycle (***North et al., 2020***). MTA salvage via the methionine salvage pathway (MSP) is also well characterised (***Sekowska and Danchin, 2002; Wray and Abeles, 1995***) (see Figure 1 B). In the classical, aerobic MSP, MTA is either processed by a two step-reaction by the MTA nucleosidase (MtnN), followed by a phosphorylation by the MTR kinase (MtnK) or by the MTA phosphorylase (MtnP). The subsequent reactions consist of a dehydration (MtnB, MTR-1P dehydratase), enolization and phosphorylation (either by MtnC or by MtnW and MtnX), deoxygenation (MtnD) and a final transamination step (MtnE) (***Sekowska et al., 2004***).

**Figure 1:**
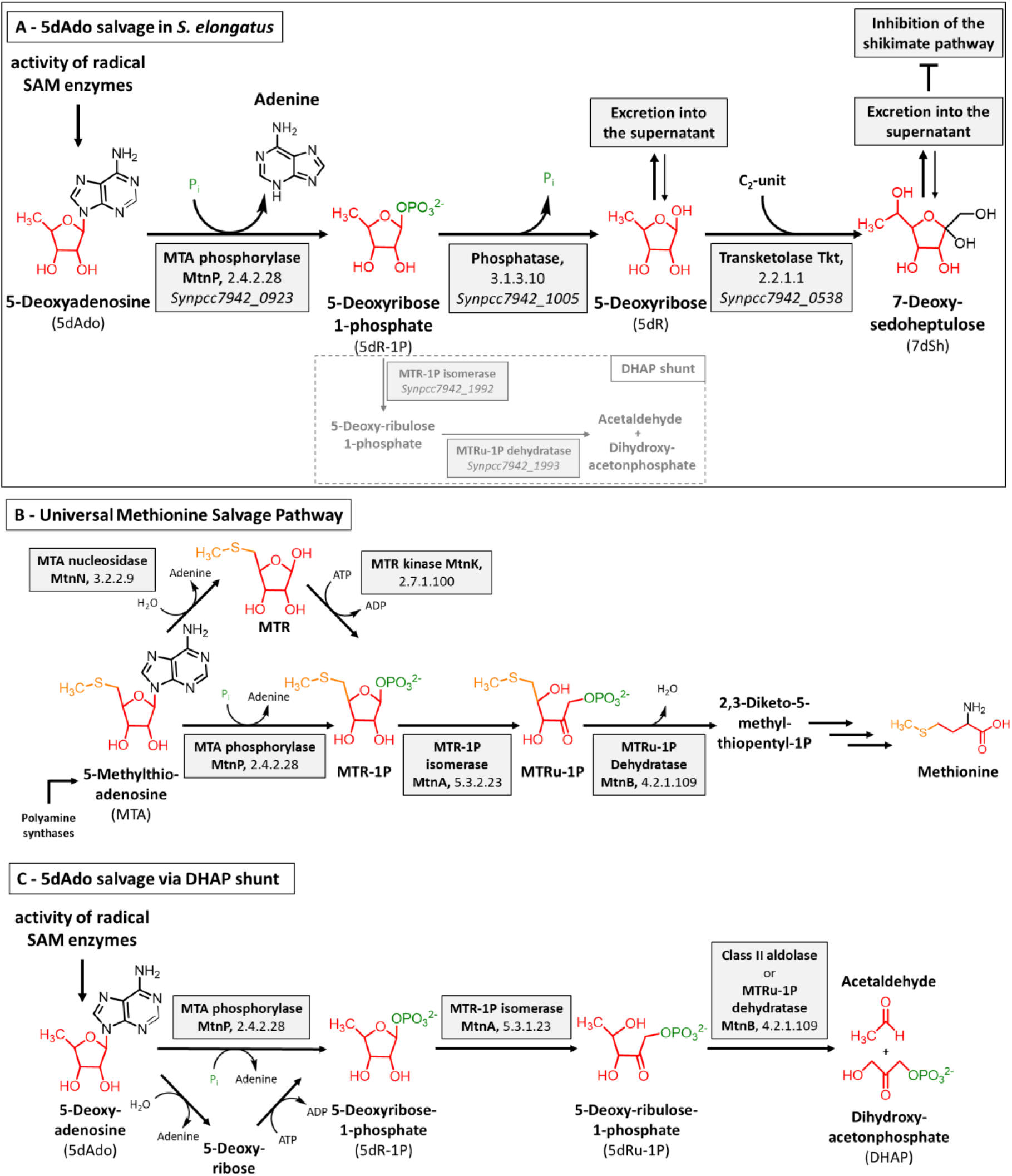
Overview of the 5dAdo and MTA salvage pathways. (A) - 5dAdo salvage in *Synechococcus elongatus* via the excretion of the bioactive deoxy-sugars 5dR and 7dSh (this study). 5dR-1P is partially also metabolized via the DHAP shunt (shown by dashed line), especially under low carbon conditions. (B) - Universal methionine salvage pathway (MSP) (***Sekowska et al., 2004***). MTR: Methylthioribose, MTRu-1P: Methylthioribulose-1P, (C) - 5dAdo salvage via the DHAP shunt (***Beaudoin et al., 2018; North et al., 2020***).

Despite the high abundance of radical SAM enzymes and thereby of 5dAdo, less is known about 5dAdo salvage. *In vitro* experiments showed that 5dAdo can be processed by a two-step reaction, in which 5dAdo is cleaved by the promiscuous MTA nucleosidase resulting in the release of adenine and 5-deoxyribose (5dR) (***Challand et al., 2009; Choi-Rhee and Cronan, 2005***). The subsequent phosphorylation of 5dR by MtnK results in the formation of 5-deoxyribose 1-phosphate (5dR-1P). The second option is the direct conversion of 5dAdo into 5dR-1P and adenine via the promiscuous MTA phosphorylase (***Savarese et al., 1981***). Therefore, Sekowska and coworkers suggested that 5dAdo salvage is paralogous to the MSP and is driven by the promiscuous activity of the enzymes of the MSP (***Sekowska et al., 2018***). Recently, a pathway for 5dR salvage was elucidated in *Bacillus thuringiensis* involving the sequential activity of a kinase (DrdK), an isomerase (DrdI) and a class II aldolase (DrdA), which are encoded by a specific gene cluster (***Beaudoin et al., 2018***). The authors propose that 5dR is phosphorylated to 5dR-1P, which is then isomerized into 5-deoxyribulose 1-phosphate (5dRu-1P) and subsequently cleaved by an aldolase into acetaldehyde and dihydroxyacetone phosphate (DHAP) for primary metabolism. In organisms that lack the specific gene cluster, the cleavage of 5dAdo into DHAP and acetaldehyde is proposed to occur via the promiscuous activity of enzymes of the MSP. In support of this hypothesis, it was shown that *Arabidopsis thaliana* DEP1, a MTR-1P dehydratase of the MSP, is promiscuous and can also cleave 5dRu-1P into DHAP and acetaldehyde, suggesting that a specific aldolase is not required for 5dAdo salvage (***Beaudoin et al., 2018***). In agreement with this, the promiscuous activity of MSP enzymes in the 5dAdo salvage was recently reported in *Methanocaldococcus jannaschii* (*M. jannaschii*) (***Miller et al., 2018***). Methylthioribose 1-phosphate isomerase (MTRI) was shown to use the substrates MTR-1P, 5dR-1P and 5dR. And only recently, it was demonstrated that 5dAdo is processed to DHAP and acetaldehyde by the first enzymes of the MSP and a clustered class II aldolase in *Rhodospirillum rubrum* and pathogenic *Escherichia coli* strains, in a process they called the “DHAP shunt” (***North et al., 2020***) (see Figure 1 C).

In our previous work, we isolated the rare deoxy-sugar – namely 7-deoxysedoheptulose (7-deoxy-D-*altro*-2-heptulose, 7dSh) – from the supernatant of the unicellular cyanobacterium *Synechococcus elongatus* PCC 7942 (henceforth referred to as *S. elongatus*) (***Brilisauer et al., 2019***). This compound showed bioactivity towards various prototrophic organisms, e.g. other cyanobacteria, especially *Anabaena variabilis* ATCC 29413 (henceforth referred to as *A. variabilis*), *Saccharomyces* and *Arabidopsis*. We hypothesized that 7dSh is an inhibitor of the enzyme dehydroquinate synthase (DHQS, EC 4.2.3.4) (***Brilisauer et al., 2019***), the second enzyme of the shikimate pathway. Because of the streamlined genome of *S. elongatus* and the lack of specific gene clusters for the synthesis of secondary metabolites (***Copeland et al., 2014; Shih et al., 2013***) the pathway for 7dSh synthesis remained enigmatic. It is worth to mention that 7dSh was also isolated from the supernatant of *Streptomyces setonensis (**Brilisauer et al., 2019; Ito et al., 1971**).* Even in this species, a pathway for 7dSh biosynthesis has remained unresolved. We speculated that 7dSh might be synthesized via primary metabolic pathways due to enzyme promiscuity. Enzyme promiscuity, the ability of an enzyme to use various substrates, is especially important for organisms with a small genome. Previously it was described that the marine cyanobacterium *Prochlorococcus* uses a single promiscuous enzyme that can transform up to 29 different ribosomally synthesized peptides into an arsenal of polycyclic bioactive products (***Li et al., 2010***). As from the 7dSh-containing supernatant of *S. elongatus* we additionally isolated the deoxy-sugar 5-deoxy-D-ribose (5dR), we hypothesized that 5dR could serve as a putative precursor molecule of 7dSh (***Brilisauer et al., 2019***). *In vitro,* 5dR can serve as a substrate for a transketolase-based reaction, in which a C_2_-unit (e.g. from hydroxypyruvate) is transferred to the C_5_-unit (3*S*, 4*R* configurated) leading to the formation of 7dSh (***Brilisauer et al., 2019***).

In this work we identified the pathway for 7dSh biosynthesis, which involves a new salvage route for 5dAdo resulting in the release of 5dR and 7dSh in the culture medium. Therefore, *S. elongatus* can synthesize an allelopathic inhibitor from the products of the primary metabolism by using promiscuous enzymes.

## Results

### 5dR and 7dSh accumulation in supernatants of *S. elongatus* is strongly promoted by CO_2_ supplementation

Previously we estimated the content of 7dSh in the supernatant of *S. elongatus* cultures via a bioassay based on the size of the inhibition zone of *A. variabilis* exposed to the supernatant of *S. elongatus (**Brilisauer et al., 2019**)*. The content of 5dR was neither estimated nor quantified before. To decipher the biosynthesis of 5dR and 7dSh in *S. elongatus*, we developed a gas chromatography-mass spectrometry (GC-MS)-based method that enables the detection and absolute quantification of low μM concentrations of these metabolites in the supernatant of cyanobacterial cultures (see Material and Methods). Briefly, 200 μL of culture supernatant were lyophilized and extracted with chloroform, methanol, and H_2_O. Subsequently, the polar phase was chemically derivatised with methoxylamine and MSTFA for GC-MS analysis. As already reported (***Brilisauer et al., 2019***), 7dSh accumulation in the supernatant requires elevated CO_2_ supply to the cultures (Figure 2 C). Under 2 % CO_2_ supplementation, 5dR gradually accumulated with increasing optical density of the cultures, whereas 7dSh accumulation only started during a later growth phase. After 30 days of growth, the amount of 7dSh in the supernatant was around one quarter compared to that of 5dR. 5dR accumulation was strongly promoted by CO_2_ supplementation, however, a small amount already started to accumulate in the cultures under ambient air conditions (Figure 2 B).

**Figure 2:**
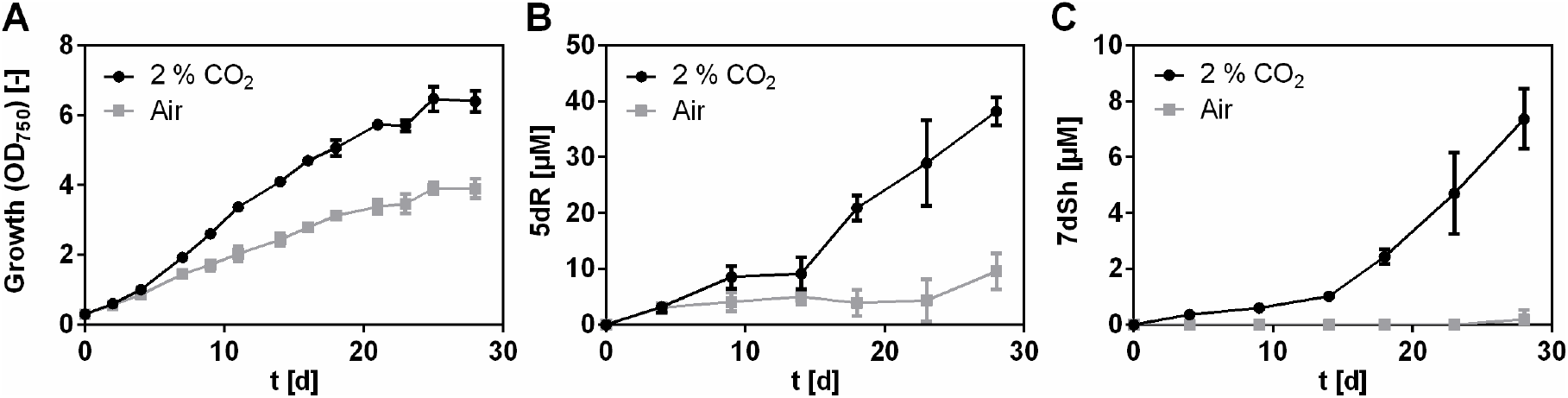
5dR and 7dSh accumulation in the supernatant of *S. elongatus* is strongly promoted by high CO_2_ concentrations. *S. elongatus* cultures aerated either with ambient air (grey squares) or with air supplemented with 2 % CO_2_ (black dots). (A) Over time growth of *S. elongatus* (indicated by OD_750_). Over time concentration of 5dR (B) or 7dSh (C) in the supernatant of *S. elongatus* cultures. Note the different values of the y-axis. Data shown represent mean and standard deviation of three independent biological replicates.

Although the optical density of the aerated cultures in the last days of the experiment reached values that were similar to those of the CO_2_-supplemented cultures, where 7dSh accumulation started, 7dSh could never be detected in air-grown cultures. This suggests that the formation of the deoxy-sugars is not only dependent on a certain cell density, but is also related to a specific metabolic state.

To gain further insights into 5dR/7dSh metabolism, we measured the intracellular concentration of 5dR and 7dSh over the whole cultivation process but were hardly able to detect any of either deoxy-sugar (Figure S1). Moreover, the small intracellular amount remained nearly constant while the extracellular concentration increased. The fact that 5dR and 7dSh only accumulate in the supernatant but is almost undetectable intracellularly strongly suggests that extracellular 5dR/7dSh accumulation is not due to cell lysis but due to secretion of the compounds immediately after their formation. Removal of these metabolites from the cytoplasm is probably essential for *S. elongatus* as both molecules showed growth inhibition towards the producer strain at elevated concentrations (Figure 3). 7dSh is bactericidal at concentrations of 100 μM, while 5dR is bacteriostatic at concentrations of 250 μM.

**Figure 3:**
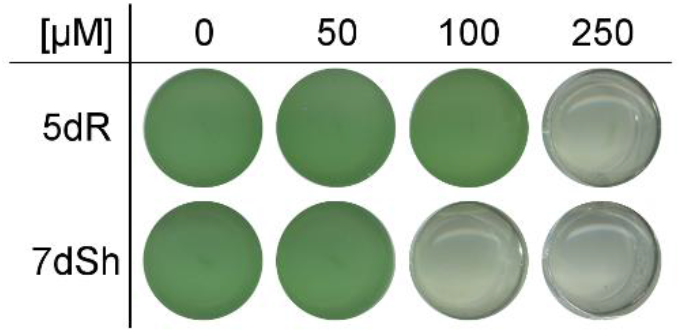
5dR and 7dSh are inhibiting the growth of the producer strain. Effect of different concentrations of 5dR and 7dSh on the growth of *S. elongatus*. The cultures were inoculated at OD_750_ = 0.1 in 1 mL BG11 medium in the absence (0) or presence of either 5dR or 7dSh at the indicated concentrations and grown in a 24-well plate for 3 days. The experiment was performed in triplicates. The results of one replicate are shown.

### 5dR is a precursor molecule for 7dSh biosynthesis *in vivo*

In our previous work, we reported the *in vitro* synthesis of 7dSh by converting 5dR into 7dSh by a transketolase-based reaction with hydroxypyruvate as a C_2_-unit donor (***Brilisauer et al., 2019***). To determine whether 5dR might also be a precursor molecule for 7dSh *in vivo*, a 5dR-feeding experiment was performed (Figure 4). To unambiguously distinguish the naturally formed and the supplemented 5dR, uniformly labelled [U-^13^C_5_]-5dR (^13^C_5_-5dR) was synthesized and added at a final concentration of 20 μM to *S. elongatus* cultures at the beginning of the cultivation. The concentration of labelled (Figure 4 B, C), unlabelled (Figure 4 D, E) and the total amount of 5dR and 7dSh (Figure 4 F, G) was determined by GC-MS at different time points over a period of 30 days.

**Figure 4:**
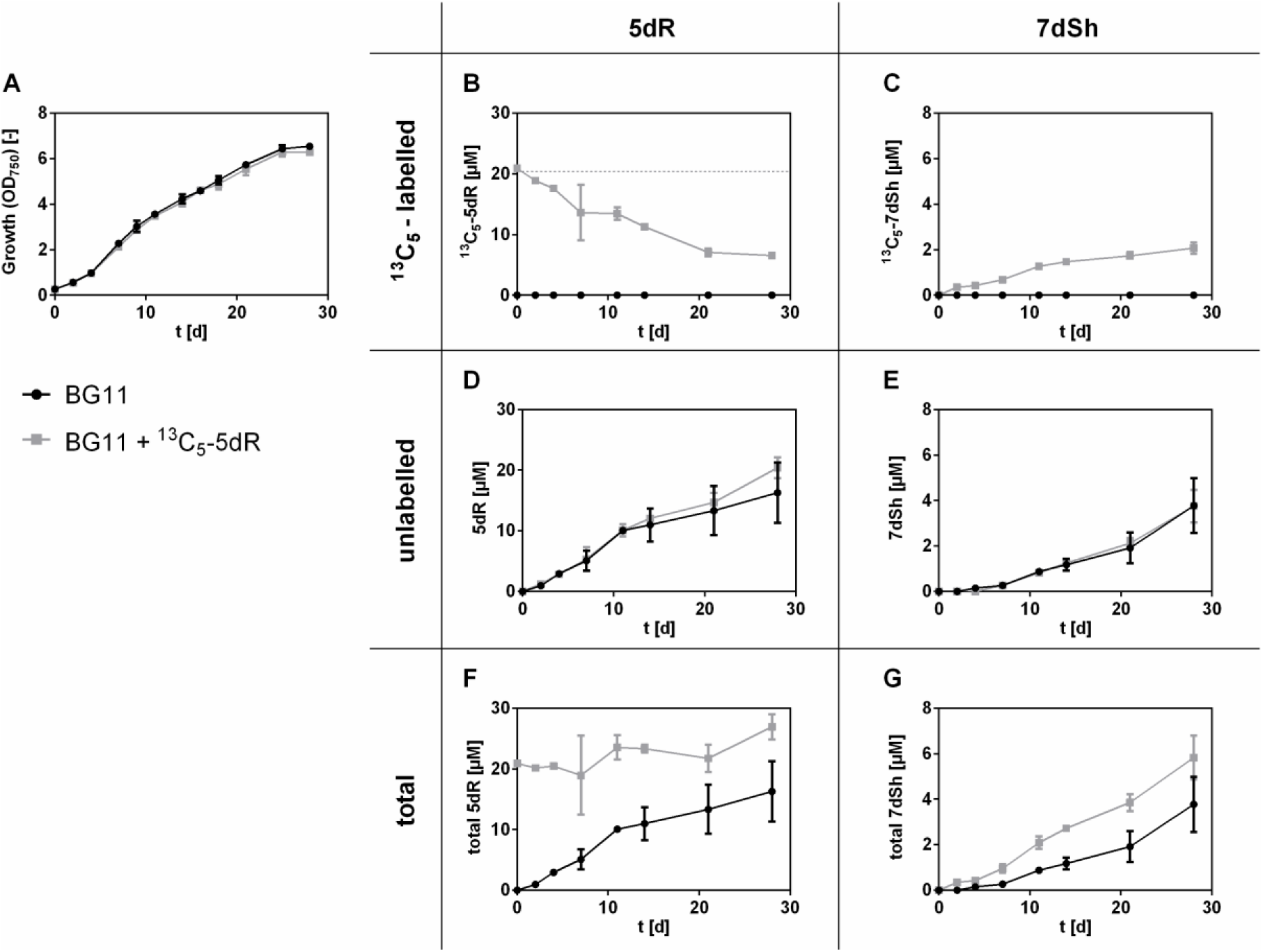
5dR is the precursor molecule of 7dSh. Effects of ^13^C_5_-5dR supplementation over the time on the growth of *S. elongatus* (A) or on the concentration of ^13^C_5_-5dR (B), ^13^C_5_-7dSh (C), unlabelled 5dR (D), unlabelled 7dSh (E), total 5dR (F) and total 7dSh (G) in the culture supernatant. 20 μM ^13^C_5_-5dR (indicated by dashed line) was added at the beginning of the cultivation (grey squares). Control cultures (black dots) were cultivated in BG11 without supplemented ^13^C_5_-5dR. All cultures were aerated with air supplemented with 2 % CO_2_. Values shown in the graphs represent mean and standard deviation of three biological replicates.

Neither the growth of *S. elongatus* nor the excretion of unlabelled, intracellular synthesized 5dR and 7dSh was affected by the addition of exogenous ^13^C_5_-5dR (Figure 4 A, D, E). We found that ^13^C_5_-5dR is taken up by the cultures as its concentration in the supernatant continuously decreased (Figure 4 B, grey squares). Already within 2 days, ^13^C_5_-7dSh could be detected in the supernatant of these cultures (Figure 4 C, grey squares), clearly proving that ^13^C_5_-7dSh was formed from the precursor molecule ^13^C_5_-5dR. However, only a small amount of exogenously added ^13^C_5_-5dR was converted into 7dSh. At the end of the experiment, 10 % of the initially applied ^13^C_5_-5dR (20 μM) was converted into ^13^C_5_-7dSh (around 2 μM). Around 30 % of ^13^C_5_-5dR remained in the supernatant (6.5 μM). The residual amount is assumed to be metabolised via (an)other pathway(s). Because unlabelled 5dR was excreted at the same time as ^13^C_5_-5dR was taken up (Figure 4 B, D), we conclude that 5dR must be imported and exported in parallel.

### 5dAdo as a precursor molecule of 7dSh

Next, we asked the question where 5dR is derived from and this drew our attention to 5dAdo, a byproduct of radical SAM enzymes (***Wang and Frey, 2007***). The compound has to be removed because of its intracellular toxicity (***Choi-Rhee and Cronan, 2005***), and its cleavage can result in the formation of 5dR (***Beaudoin et al., 2018; Choi-Rhee and Cronan, 2005***) (see Figure 1 C). To prove the hypothesis that 7dSh is formed as a result of 5dAdo salvage in *S. elongatus*, 5dAdo feeding experiments were performed and the supernatants were analysed by GC-MS (Figure 5). Notably, the growth of *S. elongatus* was not affected by supplementation with 5dAdo, which was taken up very quickly (Figure 5 A, B). After 4 days, almost all 5dAdo was taken up. A control experiment showed that the rapid decline in the amount of 5dAdo in the supernatant was not caused by the instability of 5dAdo in the medium. Feeding of the cells with 5dAdo immediately led to an enhanced accumulation of 5dR in the culture supernatant (Figure 5 C). After 14 days, 7dSh accumulation in 5dAdo-supplemented cultures was clearly enhanced in comparison to control cultures (Figure 5 D), supporting our hypothesis that 5dAdo is a precursor molecule of 7dSh. This experiment also revealed that only about half of the supplemented 5dAdo (initial concentration: 25 μM) is converted into 5dR and 7dSh, because at the end of the growth experiment the 5dR content in the supplemented cultures is increased by around 10 μM, and that of 7dSh by 2 μM. This suggest that other pathway(s) for 5dAdo salvage must exist.

**Figure 5:**
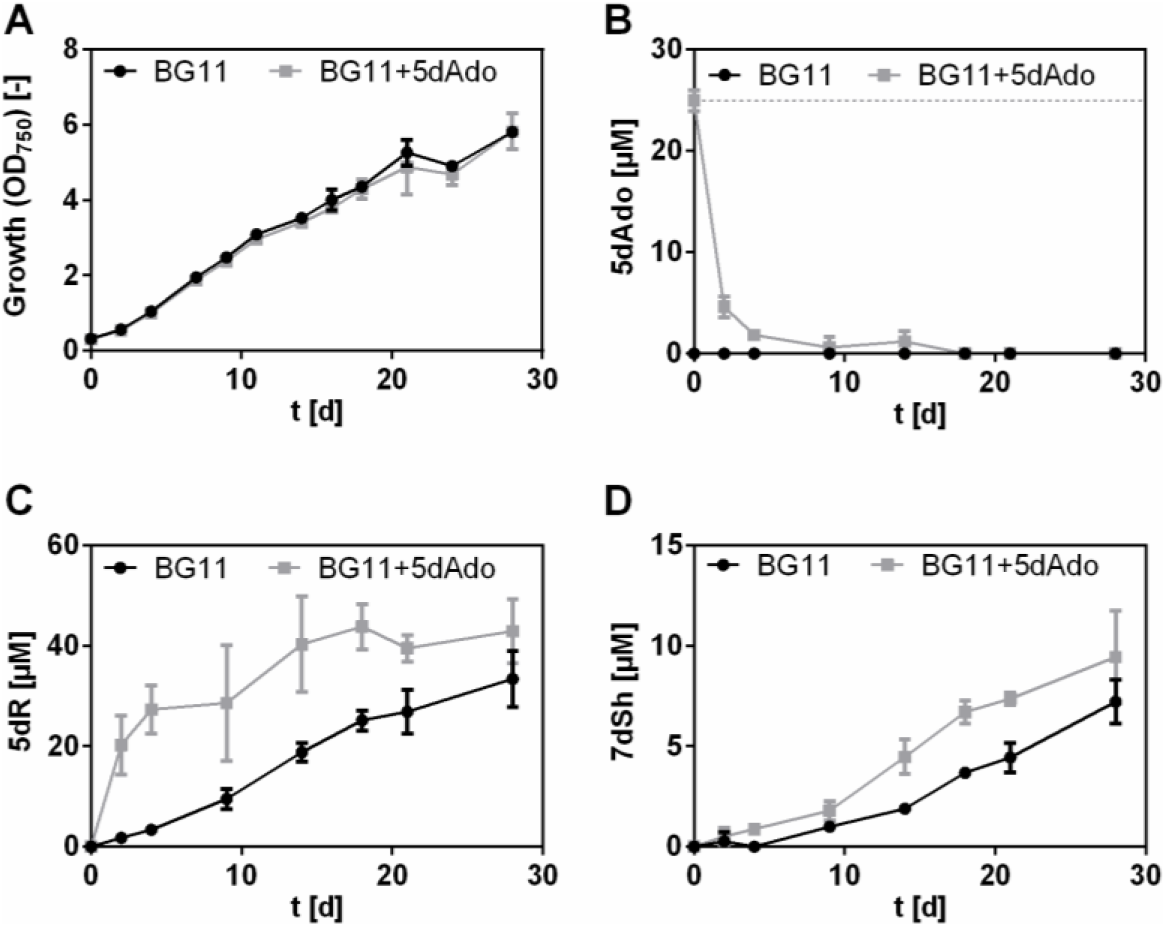
5dAdo feeding experiment. Effect of 5dAdo-supplementation on the growth of *S. elongatus* (A) or on the concentration of 5dAdo (B), 5dR (C) and 7dSh (D) in the culture supernatant. 25 μM 5dAdo (indicated by dashed line) was added at the beginning of the cultivation (grey squares). Control cultures (black dots) were cultivated in BG11 in absence of exogenous 5dAdo. All cultures were aerated with air supplemented with 2 % CO_2_. Note the different values of the y-axis. Values shown in the graphs represent mean and standard deviation of three biological replicates.

5dAdo is known to be cleaved either by the MTA nucleosidase (MtnN) or by the MTA phosphorylase (MtnP) (***Challand et al., 2009; Choi-Rhee and Cronan, 2005; Savarese et al., 1981***). The first reaction leads to the release of adenine and 5dR, the latter is phosphate-dependent and leads to the release of adenine and 5dR-1P. In *S. elongatus,* no homologous gene for a MTA nucleosidase was found, but gene *Synpcc7942_0923* is annotated as a MTA phosphorylase. Therefore, an insertion mutant of *Synpcc7942_0923* was generated via the replacement of the gene by an antibiotic resistance cassette (*S. elongatus mtnP*::*spec_R_*). Under conditions favourable for 5dR/7dSh production the mutant grew like the wild type (Figure 6 A). A GC-MS analysis of the culture supernatant revealed that the mutant neither excreted 5dR nor 7dSh (Figure 6 C, D). Instead, while undetectable in the supernatant of the wild type strain, 5dAdo strongly accumulated in the supernatant of *S. elongatus mtnP*::*spec_R_* cultures (Figure 6 B). This clearly showed that 5dR/7dSh are derived from 5dAdo in an MtnP dependent manner. Due to the detoxification via excretion, the *mtnP*::*spec_R_* mutant escapes the toxic effect of 5dAdo and does not show any growth disadvantage (Figure 6 A). It has previously been reported that a *mtnP* knockout mutant in *S. cerevisiae* as well as MtnP-deficient mammalian tumour cells excreted MTA (***Chattopadhyay et al., 2006; Kamatani and Carson, 1980***). Both MTA and 5dAdo are known to be cleaved by MtnP (***Savarese et al., 1981***). Consistently, the *mtnP*::*spec_R_* mutant excretes MTA as well as 5dAdo (Figure 6 E). As 5dR/7dSh formation is strongly dependent on the cultivation at elevated CO_2_ concentration we measured the amount of 5dAdo and MTA in cultures of the *mtnP*::*spec_R_* mutant supplied with ambient air or with air enriched with 2 % CO_2_. It turned out that the amounts of excreted 5dAdo and MTA (normalized to the optical density of the cultures) are almost identical under atmospheric or elevated CO_2_ conditions (Figure 6 E). This clearly indicates that 5dAdo salvage via 5dR/7dSh formation and excretion at high CO_2_ conditions is not triggered by an increased synthesis of the precursor molecule 5dAdo compared to ambient CO_2_ concentrations. Rather, it appears that 5dAdo is actively metabolised into 5dR/7dSh under elevated CO_2_ conditions, whereas 5dAdo salvage under ambient CO_2_ conditions is conducted by (an)other pathway(s). Since the MTA formation is also unaltered (Figure 6 E), we exclude that 5dAdo salvage via 5dR/7dSh formation is triggered by an enhanced demand of MTA salvage via a bifunctional MTA/5dAdo salvage pathway.

**Figure 6:**
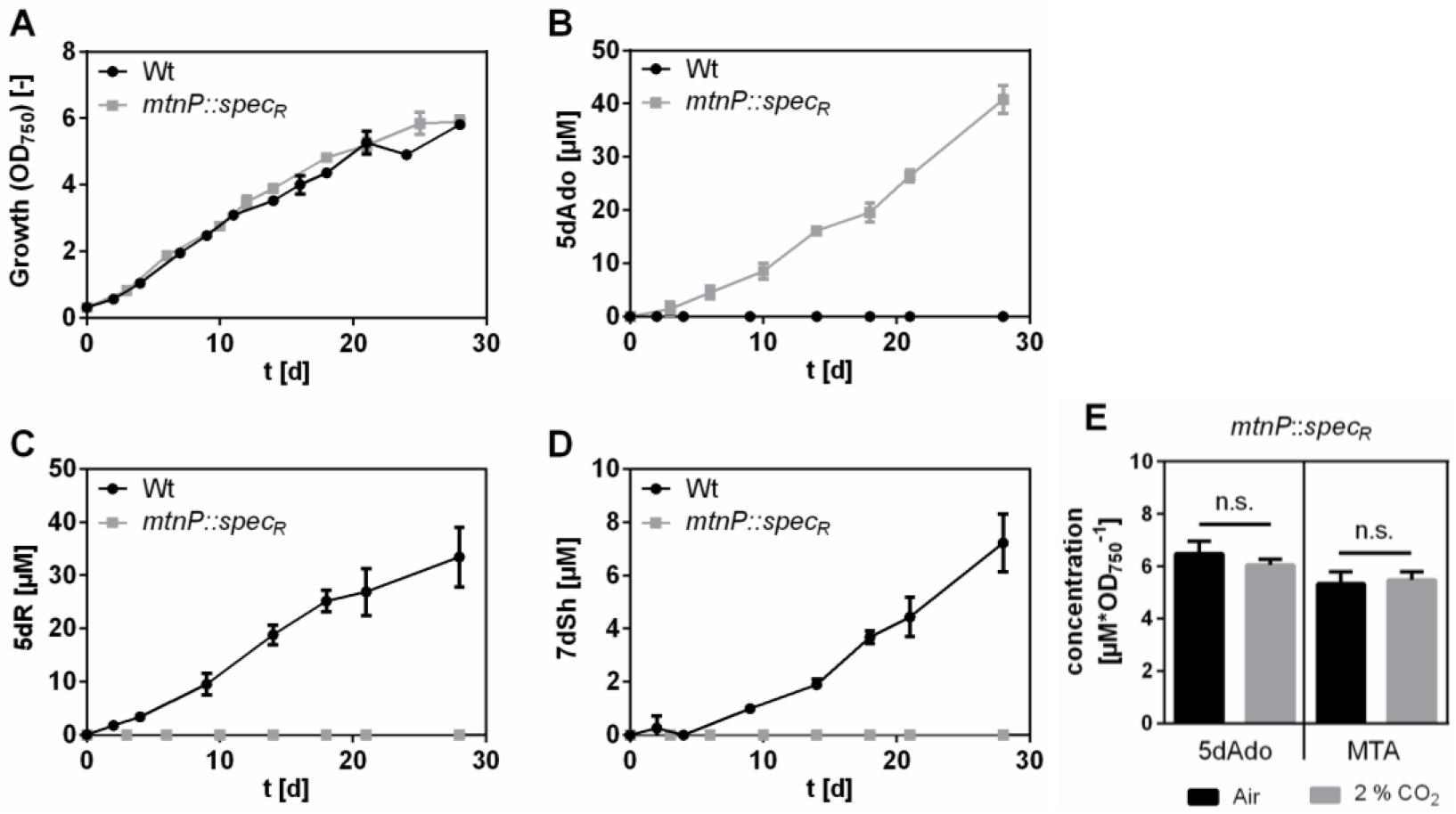
5dAdo is cleaved by MtnP and then metabolised into 5dR and 7dSh in *S. elongatus* at high CO_2_ concentrations. Growth (A), concentrations of 5dAdo (B), 5dR (C) and 7dSh (D) in the supernatant of *S. elongatus* wild type (black dots) or *mtnP*::*spec_R_* mutant (grey squares). All cultures were aerated with air supplemented with 2 %CO_2_. Note the different values of the y-axis. (E) 5dAdo and MTA concentrations in the supernatant of the *mtnP*::*spec_R_* mutant normalised on the optical density after 11 days of cultivation (cultures were either aerated with atmospheric air (black) or with air supplemented with 2 % CO_2_ (grey)). Values shown in the graphs represent mean and standard deviation of three biological replicates.

### 5dR and 7dSh formation is not ubiquitous

To clarify how widespread the synthesis of 7dSh or 5dR is in cyanobacteria, we analysed the supernatants of other cyanobacterial strains via GC-MS (*Synechococcus* sp. PCC 6301, *Synechococcus* sp. PCC 7002, *Synechococcus* sp. PCC 6312, *Synechococcus* sp. PCC 7502, *Synechocystis* sp. PCC 6803, *Anabaena variabilis* ATCC 29413, *Nostoc punctiforme* ATCC 29133, *Anabaena* sp. PCC 7120). Only in three of five *Synechococcus* strains, the deoxy-sugars 5dR and 7dSh were detectable in the supernatant. All the other strains accumulated neither 5dR nor 7dSh. In the freshwater strain *Synechococcus* sp. PCC 6301, the amounts of 7dSh and 5dR were in a similar concentration range to those in *S. elongatus*. This is not surprising since the genome of *Synechococcus* sp. PCC 6301 is nearly identical to that of *S. elongatus* PCC 7942 (***Sugita et al., 2007***). Very small amounts of 5dR and 7dSh were detected in the marine strain *Synechococcus* sp. PCC 7002. In *Streptomyces setonensis*, which was shown to produce 7dSh (***Brilisauer et al., 2019; Ito et al., 1971***), we detected 113 ± 7 μM 7dSh but no 5dR in the supernatant of cultures grown for 7 days.

### 5dAdo cleavage is strictly dependent on phosphorylase activity

To further characterize the cleavage of 5dAdo in *S. elongatus*, a crude extract assay was performed. Crude extracts of *S. elongatus* wild type and *mtnP*::*spec_R_* mutant cells were incubated with 5dAdo in the presence or absence of potassium phosphate buffer (PPB). Analysis of the extracts via thin layer chromatography revealed that 5dAdo cleavage and, thereby, adenine release is strictly dependent on the presence of phosphate (Figure 7). Adenine is only released by *S. elongatus* cell extracts in the presence of potassium phosphate buffer (red label), but not by *mtnP*::*spec_R_* mutant cells, which are not capable of 5dAdo cleavage. Therefore, 5dAdo cleavage in *S. elongatus* is strictly dependent on the presence of the MTA phosphorylase. Other enzymes, for example purine nucleosidase phosphorylases (***Lee et al., 2004***), apparently do not process 5dAdo in the *S. elongatus* cell extract. This result implies that the first product of 5dAdo cleavage must be 5dR-1P, which is subsequently converted into 5dR. 5dR-1P seemed quite stable because liquid chromatography (LC)-MS analysis revealed that a compound with a *m/z* ratio that corresponds to the sum formula of 5dR-1P ([M+H, M+Na]+ (*m/z* 215.0315; 237.0135)) accumulated in the crude extract (Figure S2). Furthermore, no 5dR formation was observed in the crude extracts (Figure S3).

**Figure 7:**
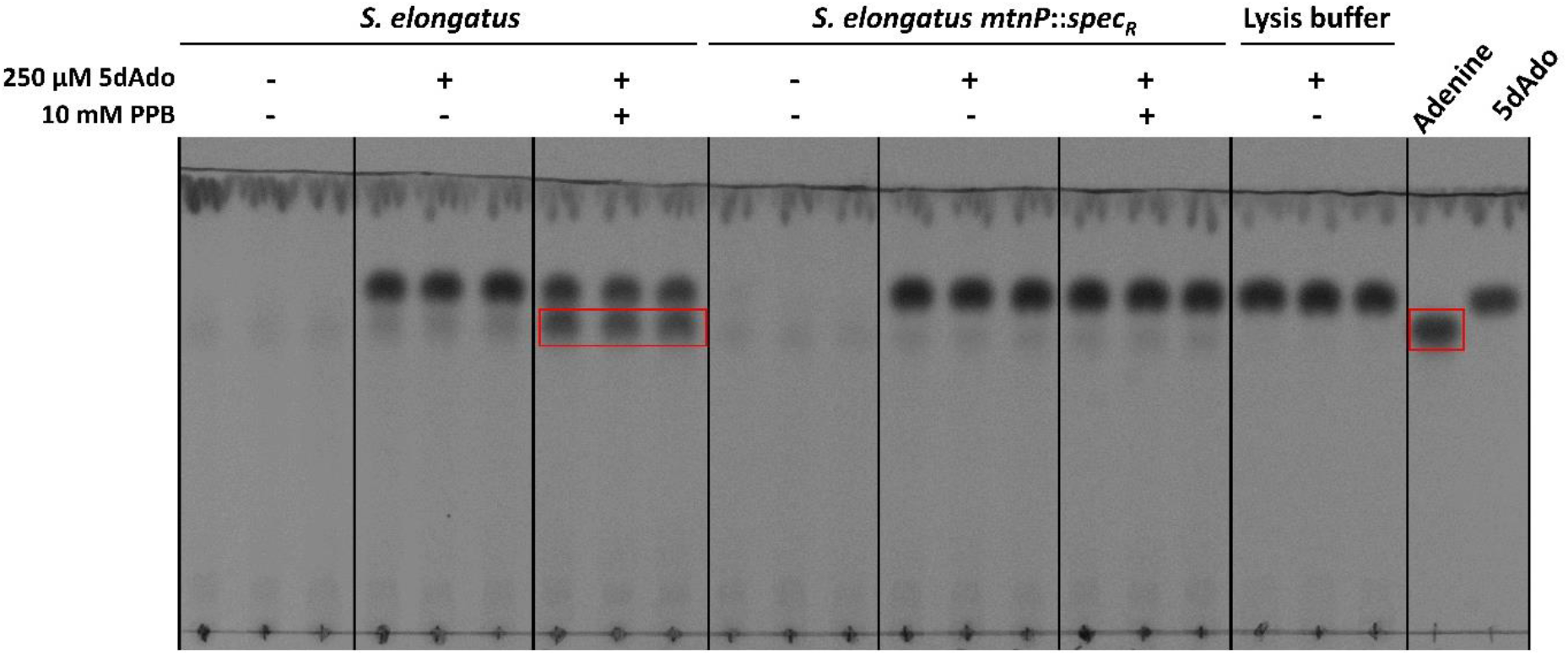
5dAdo cleavage in *S. elongatus* is phosphate dependent. Crude extracts from *S. elongatus* or *S. elongatus mtnP*::s*pec_R_* were incubated with 5dAdo in the presence or absence of potassium phosphate buffer (PPB) and then analysed via thin layer chromatography on silica gel. 5dAdo (R_f_=0.68) and adenine (R_f_=0.76) analytes were visualized via absorption at 254 nm. Pure adenine and 5dAdo were used as standards (right). Spots corresponding to adenine are highlighted with a red box. Three independent replicates are shown for each condition. The stability of 5dAdo in the buffer is shown with the lysis buffer control.

With this, we exclude a spontaneous hydrolysis of 5dR-1P which is in accordance to the literature, where 5dR-1P is reported to be metabolically stable (***Plagemann and Wohlhueter, 1983***).

### 5dR-1P is dephosphorylated by a specific phosphatase

As 5dR-1P is metabolically stable we assumed the involvement of a specific phosphatase in 5dR-1P dephosphorylation. The dephosphorylation of 5dR-1P in 5dAdo salvage is surprising as in the literature it is suggested that the phosphorylation of 5dR is essential for its further metabolization (***Beaudoin et al., 2018; North et al., 2020; Sekowska et al., 2018***). To identify the responsible phosphatase, we analysed the genome of *S. elongatus* regarding the presence of phosphoric monoester hydrolases (see Table S3, Supporting information). *Synpcc7942_1005*, which is annotated as glucose-1-phosphatase, belonging to the haloacid dehalogenase (HAD)-like hydrolase superfamily subfamily IA (***Burroughs et al., 2006; Koonin and Tatusov, 1994***), seemed a promising candidate as only *S. elongatus* and *Synechococcus* sp. PCC 6301, which both produce larger amounts of 5dR/7dSh, possess a homologous gene. The other cyanobacteria mentioned above do not possess it. Furthermore, phosphatases from the HAD-like hydrolase superfamily are known to be promiscuous enzymes dephosphorylating various phosphate-sugars (***Kuznetsova et al., 2006; Pradel and Boquet, 1988***). To examine whether this gene is essential for 5dR-1P dephosphorylation or 5dR/7dSh synthesis, an insertion mutant of this gene was created by the replacement with a spectinomycin resistance cassette (*S. elongatus Synpcc7942*_*1005*::*spec_R_*). Under 5dR/7dSh production conditions (air supplemented with 2 % CO_2_) the mutant grew like the wildtype (Figure 8 A). Whereas the wildtype excretes 5dR and 7dSh, the mutant only excretes minor amounts of 5dR and no 7dSh (Figure 8 C, D). Instead, the mutant excreted 5dAdo, which was never detected in the supernatant of the wildtype (Figure 8 B). This clearly shows that the gene product of *Synpcc7942_1005* is the major enzyme for the dephosphorylation of 5dR-1P. However, since even in the mutant small quantities of 5dR were detectable, we suggest that in addition to the gene product of *Synpcc7942_1005* other phosphatases may contribute to residual 5dR-1P dephosphorylation. In agreement with this conclusion, we found that *Synechoccoccus* sp. PCC 7002, which does not possess a homolog of *Synpcc7942_1005* also excreted minor amounts of 5dR and 7dSh (see above).

**Figure 8:**
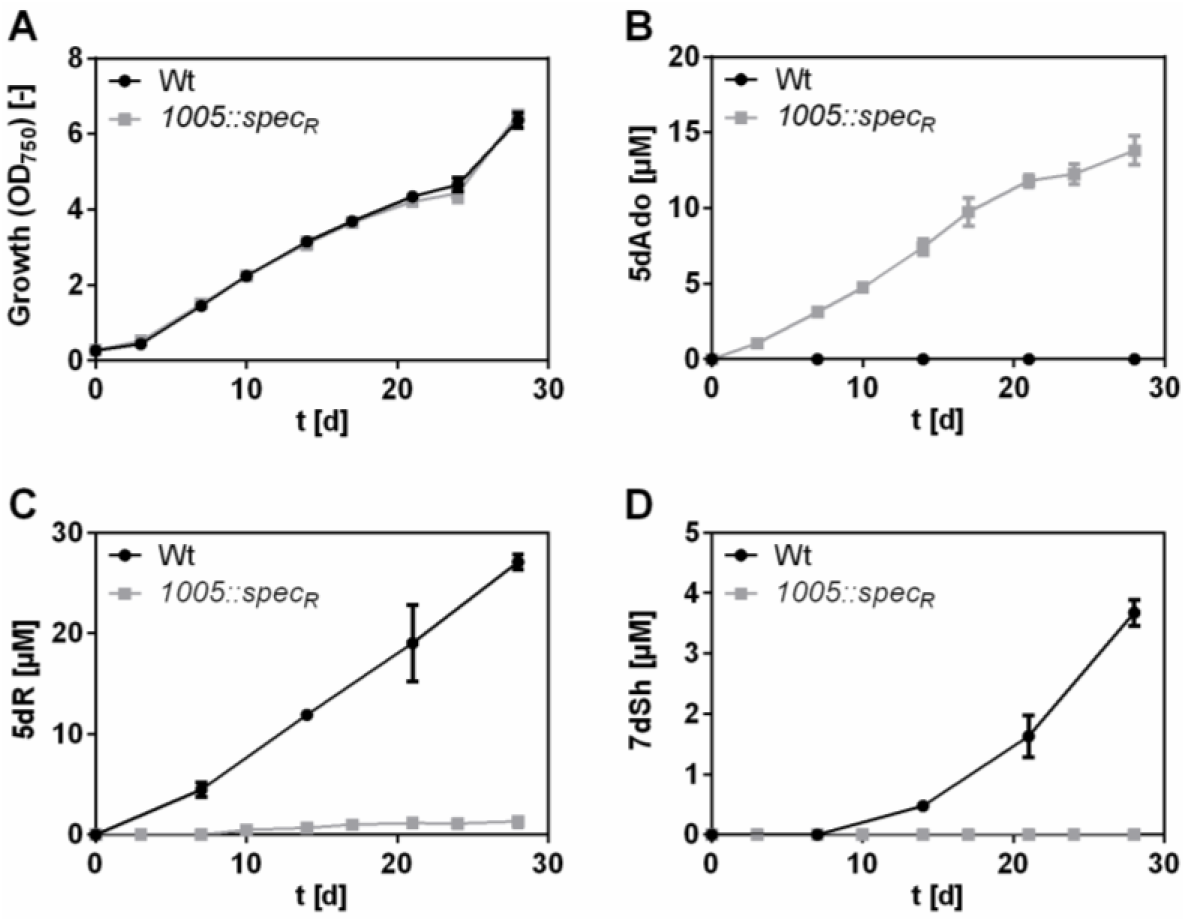
5dR-1P is dephosphorylated by a phosphatase from the HAD hydrolase superfamily (*Synpcc7942_1005*, EC 3.1.3.10). Growth (A), concentrations of 5dAdo (B), 5dR (C) and 7dSh (D) in the supernatant of *S. elongatus* wild type (black dots) or *1005*::*spec_R_* mutant (grey squares). All cultures were aerated with air supplemented with 2 %CO_2_. Note the different values of the y-axis. Values shown in the graphs represent mean and standard deviation of three biological replicates.

### 5dR/7dSh producers possess complete MSP gene clusters

By analysing the genomes of all examined cyanobacteria in this study, it turned out that those strains that do not produce 5dR and 7dSh only possess annotated genes for the first two reactions of the MSP (*mtnP* and *mtnA*), whereas the producer strains possess annotated genes for the whole MSP pathway (Table S1). The 5dAdo salvage via the DHAP shunt requires a specific class II aldolase, e.g. DrdA in *B. thuringiensis* (***Beaudoin et al., 2018***) or Ald2 in *R. rubrum* (***North et al., 2020***), which is clustered with the first enzymes of the MSP, e.g. MtnP or MtnN/MtnK and MtnA in *R. rubrum*) or with a specific phosphorylase and isomerase as shown for *B. thuringiensis. In vitro* data suggest that the MTRu-1P-dehydratase (MtnB) from the MSP can also act as a promiscuous aldolase thereby completing the 5dAdo salvage via the DHAP shunt (***Beaudoin et al., 2018; North et al., 2020***). Since none of the analysed strains possesses such an Ald2 homolog, we assume that 5dR/7dSh producing strains might use the DHAP shunt by using MtnB under certain conditions. The 5dR/7dSh non-producer strains must employ another pathway of 5dAdo salvage or use another aldolase for the DHAP shunt.

## Discussion

Radical SAM enzymes are important enzymes in all domains of life (***Sofia et al., 2001***). A byproduct of the activity of these enzymes is 5dAdo (***Wang and Frey, 2007***). Its accumulation inhibits the activity of the radical SAM enzymes themselves (***Challand et al., 2009; Choi-Rhee and Cronan, 2005; Farrar et al., 2010; Palmer and Downs, 2013***). Therefore, 5dAdo salvage pathways are essential. In this study we showed that the unicellular cyanobacterium *S. elongatus* PCC 7942 has a special salvage route for 5dAdo, which was never reported before (Figure 1 A). We show that 5dAdo salvage can be achieved by the excretion of 5-deoxyribose and 7-deoxysedoheptulose. 5dR as a product of 5dAdo cleavage was postulated (***Parveen and Cornell, 2011; Sekowska et al., 2018***) or observed before but only in *in vitro* assays (***Choi-Rhee and Cronan, 2005; Sekowska et al., 2018***). Beaudoin and coworkers suggested 5dR excretion as a detoxification strategy for organisms that do not possess a specific gene cluster for 5dAdo salvage (***Beaudoin et al., 2018***) (analogous to MTR excretion in *E. coli* which does not possess a complete MSP (***Hughes, 2006; Schroeder et al., 1972***)). Therefore, 5dR accumulation in the supernatant of *S. elongatus* as an *in vivo* phenomenon was first reported by our previous publication (***Brilisauer et al., 2019***), and here identified as a result of 5dAdo salvage.

We propose the following model for a possible 5dAdo salvage route in *S. elongatus* by the activity of promiscuous enzymes leading to the synthesis of the bioactive deoxy-sugars 5dR and 7dSh (Figure 1 A). In brief, 5dAdo is processed by the promiscuous MTA phosphorylase into 5dR-1P. Under elevated CO_2_ conditions, this molecule is dephosphorylated to 5dR by a potentially promiscuous phosphatase to 5dR, part of which is excreted and part of which is further metabolized by the activity of a promiscuous transketolase to 7dSh, which is also excreted from the cells to avoid its inhibitory effects on the shikimate pathway (***Brilisauer et al., 2019***). 7dSh is a potent inhibitor of the dehydroquinate synthase, but the inhibitory effect of the compound is dependent on the organism. The producer strain tolerates high concentrations of 7dSh (Figure 3), whereas f.e. *A. variabilis* is highly sensitive towards 7dSh treatment (***Brilisauer et al., 2019***), suggesting that 7dSh is a potent allelopathic inhibitor.

Although most bacteria possess the enzymes for a two-step reaction of 5dAdo cleavage (MTA nucleoside and MTR kinase) (***Albers, 2009; Zappia et al., 1988***), all examined cyanobacteria possess a MTA phosphorylase (MtnP) (Table S1), which is normally present in eukaryotes (except for plants). The phenotype of the insertion mutant (*mtnP*::*spec_R_*), which excretes 5dAdo instead of 5dR/7dSh, demonstrates that 5dR and 7dSh are products of 5dAdo salvage (Figure 6). The 5dAdo salvage routes previously reported suggest that the phosphorylation of 5dR or the 5dR moiety of 5dAdo is essential to further metabolise the molecules via specific enzymes or by promiscuous activity of the enzymes of the MSP (***Beaudoin et al., 2018; North et al., 2020; Sekowska et al., 2018***). In *S. elongatus* however, 5dR-1P is subsequently dephosphorylated to 5dR for excretion or for further processing to 7dSh. Our data imply that the dephosphorylation of 5dR-1P is not due to spontaneous hydrolysis but is mainly conducted by the gene product of *Synpcc7942*_*1005* (see Figure 8). *Synpcc7942*_*1005* belongs to Mg^2+^-dependent class IA HAD-like hydrolase superfamily (***Burroughs et al., 2006***) and is annotated as a glucose-1-phosphatase, which catalyses the dephosphorylation of glucose 1-phosphate (***Turner and Turner, 1960***). As these phosphatases can also exhibit phytase activity (***Herter et al., 2006; Suleimanova et al., 2015***) we assume that the gene product of *Synpcc7942*_*1005* might also exhibits promiscuous activity, including 5dR-1P dephosphorylation. The dephosphorylation of a similar molecule (5-fluoro-5-deoxyribose 1-phosphate) by a specific phosphoesterase (FdrA) is also conducted by *Streptomyces* sp. MA37 during the production of a specific secondary fluorometabolite (***Ma et al., 2015***) (pathway shown in Figure S4).

In later growth phases, small amounts of 5dR are transformed into 7dSh which is then also immediately excreted into the supernatant (Figure 2 C, Figure 4 C, E). In our previous work we showed that the affinity of *S. elongatus* transketolase for 5dR (k_m_=108.3 mM) is 100-fold lower than for the natural substrate D-ribose-5-phosphate (k_m_=0.75 mM) (***Brilisauer et al., 2019***). This is in accordance with the fact that 7dSh is only formed when relatively high extracellular 5dR concentrations are reached (either in later growth phases or due to the addition of externally added 5dR; note that 5dR is continuously imported and exported). Furthermore, only one tenth of ^13^C_5_-5dR is converted into ^13^C_5_-7dSh. 7dSh formation from 5dR is therefore an impressive example how a more potent “derivative” (7dSh) is formed by promiscuous enzyme activity. Interestingly, a promiscuous transketolase reaction was also suggested in later steps of anaerobic 5dAdo salvage in *M. jannaschii*, in which 5dRu-1P is cleaved into lactaldehyde and methylglyoxal (***Miller et al., 2018***). *Streptomyces setonensis* (not yet sequenced) accumulates much higher concentrations of 7dSh in the supernatant than *S. elongatus* but no 5dR at all, which supports the hypothesis that 7dSh might be derived from complete conversion of 5dR by a more specific transketolase.

In high concentrations, 5dR exhibited toxicity towards the producer strain (Figure 3). 5dR toxicity was also reported in *B. thuringiensis* (***Beaudoin et al., 2018***), but the intracellular target is not yet known. Therefore, *S. elongatus* has to steadily excrete 5dR into the supernatant to avoid intracellular toxicity. Because ^13^C_5_-5dR is taken up at the same time as unlabelled 5dR is excreted (Figure 4 B, D), it is obvious that 5dR is continuously imported and exported so that hardly any 5dR accumulates intracellularly (see Figure S1). We also assume that 7dSh is im- and exported, too. This suggests the presence of an effective export system which is essential for the survival of the producer strain.

5dAdo salvage via 5dR and 7dSh excretion was only observed when cultures were aerated with air supplemented with 2 % CO_2_ (Figure 2 B, C). Since equal amounts of 5dAdo were formed under ambient CO_2_ as under high CO_2_ conditions (Figure 6 E), we assumed that under ambient CO_2_ conditions 5dAdo salvage is conducted via (an)other pathway(s). The occurrence of (an) additional 5dAdo salvage pathway(s) in *S. elongatus* is underlined by the fact that 5dAdo is not completely metabolised into 5dR/7dSh even under high CO_2_ conditions (Figure 5). Because *S. elongatus* and the other 5dR/7dSh producers are equipped with the enzymes for the whole MSP (see Table S1), we hypothesize that 5dAdo can be also metabolised via promiscuous activity of the enzymes of the MSP via the “DHAP-shunt” resulting in the formation of DHAP and acetaldehyde (see Figure 1 A, C) as suggested for organisms that do not possess a specific gene cluster for 5dAdo salvage (***Beaudoin et al., 2018; North et al., 2020; Sekowska et al., 2018***). The formation of MTA, the starting molecule of the MSP, is almost identical under atmospheric and high carbon conditions (Figure 6 E). This indicates that 5dAdo salvage via 5dR/7dSh excretion under high CO_2_ conditions is not triggered by an increased demand of MTA salvage. It is known that intracellular CO_2_/HCO_3_^−^(Ci) exhibits regulatory functions at the metabolic and transcriptomic level (***Blombach and Takors, 2015***): CO_2_/HCO_3_^−^can alter physiochemical enzyme properties and it is known to regulate virulence and toxin production in pathogens, e.g. in *Vibrio cholerae* (***Abuaita and Withey, 2009***). In particular, cyanobacteria strongly respond to the ambient Ci supply by a multitude of metabolic adaptations such as carbon concentrating mechanisms (***Burnap et al., 2015***) and the synthesis of cAMP (***Selim et al., 2018***). As we hypothesize that the fate of 5dAdo is a regulated process, we assume that the dephosphorylation of 5dR and the subsequent formation of 7dSh molecules is not an “accident”. They are rather purposely formed metabolites, which however derive from toxic byproducts of the primary metabolism. The regulation how 5dAdo is directed towards 5dR/7dSh formation has to be further investigated.

With 18 radical SAM enzymes (see Table S2), *S. elongatus* only possesses a relatively small number of radical SAM enzymes compared to other prokaryotes (*B. thuringiensis*: 15; other Firmicutes: more than 40 (***Beaudoin et al., 2018***), *R. rubrum*: 25, *M. jannaschii*: 30 (***North et al., 2020***)). Probably the most important radical SAM enzymes under the cultivation conditions applied here are involved in cofactor biosynthesis: Lipoic acid synthase (LipA), biotin synthase (BioB) and GTP 3’,8-cyclase (MoaA), which is involved in molybdopterin biosynthesis. These cofactors are presumably equally important under atmospheric or high carbon conditions resulting in the unaltered 5dAdo formation, thereby explaining the unaltered rate of 5dAdo formation.

7dSh can inhibit the growth of other cyanobacteria but also of plants, and was therefore suggested to be an allelopathic inhibitor by inhibiting the dehydroquinate synthase, the second enzyme of the shikimate pathway (***Brilisauer et al., 2019***). Additionally, 5dR is toxic for various organisms (Figure 3, (***Beaudoin et al., 2018***)). Despite the low concentrations of 5dR/7dSh observed under laboratory conditions it is imaginable that excretion of 5dR and 7dSh plays a role in protecting the ecological niche of the producer strains. 7dSh is a more potent inhibitor for example for *A. variabilis* than for the producer strain. A bactericidal effect for *A. variabilis* was observed at concentrations of 13 μM 7dSh (***Brilisauer et al., 2019***), whereas *S. elongatus* is affected by 100 μM (see Figure 3). In its natural environment, *S. elongatus* is able to form biofilms (***Golden, 2019; Yang et al., 2018***). In biofilms cyanobacteria tend to excrete exopolysaccharides (***Rossi and Philippis, 2015***) which can be used as a carbon source by heterotrophic members of the microbial community thereby causing locally elevated CO_2_ concentrations. This could lead to a local enrichment of 5dR and 7dSh, thereby providing a growth advantage to the producer strains protecting their niches against competing microalgae.

## Conclusion

5dAdo salvage is a less noticeable and overlooked research topic in comparison to methionine salvage from MTA. Hence, it should be further investigated above all because 5dAdo is present in all domains of life whereas MTA is only produced by specific organisms. It is possible that additional metabolites, apart from 7dSh, are derived from 5dAdo salvage in other organisms. This study shows that enzyme promiscuity is especially important for organisms with a small genome, since it enables them to produce special metabolites in absence of *ad hoc* biosynthetic gene clusters.

## Materials and Methods

### Cultivation

*Synechococcus elongatus* PCC 7942 was cultivated under photoautotrophic conditions in BG11 medium (***Rippka et al., 1979***) supplemented with 5 mM NaHCO3. Precultures were cultivated in shaking flasks at 30-50 μE at 125 rpm (27 °C). Main cultures were cultivated in 500-700 mL BG11 at 27 °C in flasks which were either aerated with air or air supplemented with 2 % CO_2_. For this purpose, cultures were inoculated with an optical density (OD_750_) of 0.2-0.5 and then cultivated for the first three days at 10 μE (Lumilux de Lux, Daylight, Osram). Later, the light intensity was set to around 30 μE. Growth was determined by measuring the optical density at 750 nm (Specord 205, Analytik Jena). For feeding experiments the cultures were supplemented at the beginning of the cultivation with 5dR, [U-^13^C_5_]-5dR or 5-deoxyadenosine (5dAdo; Carbosynth Ltd.) at the respective concentrations (see Results section). The other cyanobacterial strains (*Synechococcus* sp. PCC 6301, *Synechococcus* sp. PCC 6312, *Synechococcus* sp. PCC 7502, *Synechocystis* sp. PCC 6803, *Anabaena variabilis* ATCC 29413, *Nostoc punctiforme* ATCC 29133, *Anabaena* sp. PCC 7120) were cultivated as described above. *Synechococcus* sp. PCC 7002 was cultivated in a 1:1 mixture of BG11 and ASN III + vitamin B_12_ (10 μg/mL) (***Rippka et al., 1979***).

*Streptomyces setonensis* SF666 was cultivated for 7 days as described in our previous work (***Brilisauer et al., 2019***).

### Chemical synthesis of 5-deoxyribose and 7-deoxysedoheptulose (7dSh)

5dR and [U-^13^C_5_]-5dR **5** were synthesized in a four-step synthesis based on literature (***Sairam et al., 2003; Zhang et al., 2013***) with additional optimization. All synthetic intermediates shown in the reaction scheme (Figure S5) were verified by thin layer chromatography (TLC), mass spectrometry and NMR. Detailed data for the ^13^C-labelled compounds are presented in the Supplementary information. The synthesis starts with the reaction of D-ribose (Sigma) or [U-^13^C_5_]-D-ribose **1** (500.1 mg, 3.22 mmol; Eurisotop) in a 4:1 mixture of acetone:methanol with SnCl_2_×2 H_2_O (1 eq) and catalytic amounts of conc. H_2_SO_4_ at 45 °C for 20 h. After cooling to room temperature, the mixture was filtered, neutralised with NaHCO_3_ solution, once again filtered and the organic solvent was evaporated. The remaining aqueous solution was extracted with ethylacetate, dried over Na_2_SO_4_ and evaporated in vacuo to yield the acetonide-protected ribose **2** as a colourless oil (399.7 mg, 1.91 mmol, 59 %).

Envisaging the following deoxygenation reaction, the protected pentose **2** (399.7 mg, 1.91 mmol) was diluted in DCM with addition of TEA (2.5 eq). After cooling on ice, mesylchloride (2.5 eq) was slowly added and then stirred for 5 h on ice. The reaction mixture was washed with 1 N HCl, ultrapure water, NaHCO_3_ solution, NaCl solution and again with ultrapure water. The organic solvent was dried over Na_2_SO_4_ and evaporated in vacuo to give **3** as a yellowish oil (556.5 mg, 1.97 mmol, 103 %, mesylchloride as impurity), which becomes crystalline at 4 °C.

For the reduction as the third step **3** (556.1 mg, 1.91 mmol, maximum educt amount) was diluted in DMSO. After cooling on ice NaBH_4_ (5 eq) was added slowly. Afterwards the reaction mixture was heated slowly to 85 °C and reacting for 12 h. After cooling on ice, 5 % AcOH was added to quench remaining NaBH_4_. The aqueous solution was extracted with DCM, washed with ultrapure water, dried over Na_2_SO_4_ and evaporated in vacuo (40 °C, 750 mbar) to get **4** as a colourless oil (357.7 mg, 1.85 mmol, 86 %).

Deprotecting to the target **5** was achieved by diluting the acetonide-protected ω-deoxy-sugar **4** (357.7 mg, 1.85 mmol) in 0.04 N H_2_SO_4_ and heating to 85 °C for 3 h. After cooling to room temperature, the reaction mixture was neutralised with NaHCO_3_ solution and evaporated by lyophilisation. The final product was first purified by MPLC (Gradient: start CHCl_3_:MeOH 10:0; end CHCl_3_:MeOH 7:3) and HPLC (Column: HiPlexCa, 85 °C, 250×10.7 mm, 1.5 mL/min, solvent: ultrapure water) to get [U-^13^C_5_]-5-deoxy-D-ribofuranose (**5**) as a colourless oil (115.7 mg, 1.12 mmol, 61 %).

7dSh or [3,4,5,6,7-^13^C_5_]-7dSh were synthesized in a transketolase based reaction wit 5dR or [U-^13^C_5_]-5dR as substrate as described in our previous publication (***Brilisauer et al., 2019***) with slight modifications: The reaction was performed in water instead of HEPES buffer, to ensure an enhanced stability of hydroxypyruvate (very instable in HEPES (***Kobori et al., 1992***)). The reaction was performed for 7 days and fresh hydroxypyruvate was added every day. Purification was done as described for 5dR.

### Construction of insertion mutants

To create an insertion mutant of the 5’-methylthioadenosine phosphorylase (EC: 2.4.2.28, MtnP, *Synpcc7942_0932*) and of the glucose 1-phosphate phosphatase (EC: 3.1.3.10, *Synpcc7942*_1005) in *S. elongatus* PCC 7942 a spectinomycin resistance cassette was introduced inside the respective gene. An integrative plasmid was constructed in *E. coli* and then transformed into *S. elongatus*. For this purpose, flanking regions on both sides of the respective gene were amplified from *S. elongatus* colonies with primers adding an overlapping fragment (46_0923_up_fw, 47_0923_up_rev and 48_Δ0923_down_fw, 49_0923_down_rev for *Synpcc7942*_*0923*::*spec_R_*; 85_1005_up_fw, 86_1005_up_rev and 87_1005_down_fw, 88_1005_down_rev for *Synpcc7942*_*1005*::*spec_R_*). The primer sequences are shown in Table S3. The spectinomycin resistance cassette was amplified with the primers 32_Spec_fw and 33_Spec_rev from a plasmid containing the resistance cassette. All PCR amplification products were introduced into a pUC19 vector cut with XbaI and PstI by using Gibson assembly (***Gibson, 2011***). The plasmid was verified by Sanger sequencing (Eurofins Genomics). The plasmid was then transformed into *S. elongatus* using natural competence. In short, *S. elongatus* cells were harvested by centrifugation, washed with 400 μL BG11 and then incubated with 1 μg DNA in the dark for 6 h (28 °C). The cells were then plated on BG11 agar plates containing 10 μg/mL spectinomycin for three days. After that, the cells were transferred to agar plates containing 20 μg/mL spectinomycin. Segregation was confirmed by colony PCR (50_0923_rev_seg, 51_0923_fw_seg for *Synpcc7942*_*0923*::*spec_R_*; 85_1005_up_fw, 88_1005_down_rev for *Synpcc7942*_*1005*::*spec_R_*). Precultures of these strains, in the following named as *S. elongatus mtnP*::*spec_R_* or *S. elongatus 1005*::*spec_R_* were cultivated in the presence of 20 μg/mL spectinomycin. Main cultures were cultivated without antibiotic.

### Quantification of metabolites in the culture supernatant via GC-MS

Culture supernatant was collected by centrifugation of 1.5 mL culture (16.000 × *g*, 10 min, 4 °C). 200 μL of the supernatant were transferred into a 2 mL reaction tube and immediately frozen on liquid nitrogen and stored at -80 °C. The supernatant was lyophilized. For intracellular measurements, the cell pellets were also frozen in liquid nitrogen. The samples were extracted with 700 μL precooled CHCl_3_/MeOH/H_2_O (1/2.5/0.5 v/v/v) as described in the literature (***Fürtauer et al., 2016***) with slight modifications. Samples were homogenized by vortexing, ultrasonic bath (Bandelin, Sonorex) treatment (10 min) and shaking (10 min, 1.000 rpm). After that, the samples were cooled on ice for 5 min and then centrifuged (10 min, 16.000 × *g*, 4 °C). The supernatant was transferred into a new reaction tube. The pellet was again extracted with 300 μL extraction solvent as described before. The supernatants were pooled and 300 μL ice cold water was added for phase separation. The samples were vortexed, incubated on ice (5 min) and then centrifuged (10 min, 16.000 × *g*, 4 °C). 900 μL of the upper, polar phase were transferred into a new 2 mL reaction tube and dried in a vacuum concentrator (Eppendorf, Concentrator plus, mode: V-AQ, 30 °C) for approximately 4.5 h. The samples were immediately closed and then derivatized as described in the literature (***Weckwerth et al., 2004***) with slight modifications. Therefore, the pellets were resolved in 60 μL methoxylamine hydrochloride (Acros Organics) in pyridine (anhydrous, Sigma-Aldrich) (20 mg/mL), homogenized by vortexing, a treatment in an ultrasonic bath (15 min, RT) and an incubation at 30 °C on a shaker (1.400 rpm) for 1.5 h. After that, 80 μL *N*-methyl-*N*-(trimethylsilyl)trifluoroacetamide (MSTFA, Macherey-Nagel) was added and the samples were incubated at 37 °C for 30 min (1.200 rpm). The samples were centrifuged (16.000 × *g*, 2 min) and 120 μL were transferred into a glass vial with micro insert. The samples were stored at room temperature for 2 h before GC-MS measurement.

GC-MS measurements were performed on a Shimadzu GC-MS TQ 8040 (Injector: AOC-20i, Sampler: AOC-20s) with a SH-Rxi-5Sil-MS column (Restek, 30 m, 0.25 mm ID, 0.25 μm). For GC measurement, the initial oven temperature was set to 60 °C for 3 min. After that the temperature was increased by 10 °C/min up to 320 °C, which was then held for 10 min. The GC-MS interface temperature was set to 280 °C, the ion source was heated to 200 °C. The carrier gas flow (helium) was 1.28 mL/min. The injection was performed in split mode 1:10. The mass spectrometer was operated in EI mode. Metabolites were detected in MRM mode. Quantification of the metabolites was performed with a calibration curve of the respective substances (5dAdo, 5dR, 7dSh, ^13^C_5_-5dR, ^13^C_5_-7dSh).

### Quantification of MTA and 5dAdo

For the quantification of MTA and 5dAdo (Figure 6 E) 25 μL of culture supernatant were mixed with 75 μL aqueous solution of 20 % MeOH (v/v) + 0.1 % (v/v) formic acid. Samples were analysed on a LC-HR-MS system (Dionex Ultimate 3000 HPLC system coupled to maXis 4G ESI-QTOF mass spectrometer). 5dAdo and MTA were separated on a C18 column with a MeOH/H_2_O gradient (10 %-100 % in 20 min). The concentration was calculated from peak areas of extracted ion chromatograms of a calibration curve of the respective standards (MTA was obtained from Cayman Chemicals).

### Crude Extract Assays

Crude extract assays were performed by harvesting *S. elongatus* or *S. elongatus mtnP*::*spec_R_* cultures which were cultivated at 2 % CO_2_ supplementation until an optical density of around OD_750_=4 (t=14 d). 10 mL of the cultures were centrifuged (3.200 × *g*, 10 min, 4 °C), the supernatant was discarded, the pellet was washed with 10 mL fresh BG11 medium, and again centrifuged. After that, the pellet was resuspended in 2.5 mL lysis buffer (25 mM HEPES pH 7.5, 50 mM KCl, 1 mM DTT) and filled into 2 mL tubes with a screw cap. 100 μL glass beads (ø=0.1-0.11 mm) was added and the cells were then disrupted at 4 °C by a FastPrep^®^-24 instrument (MP Biomedicals, 5 m/s, 20 sec, 3x with 5 min break). To remove the cell debris a centrifugation step was performed (25.000 × *g*, 10 min, 4 °C). 200 μL of the supernatant was used for the crude extract assay. The extract was either used alone or supplemented with 10 μL 5 mM 5dAdo (final concentration: 250 μM) or in combination with 40 μl 50 mM potassium phosphate buffer (PBB) pH 7.5 (final concentration: 10 mM). The extracts were incubated at 28 °C for 7 h, then frozen in liquid nitrogen and lyophilized. 100 μL MeOH was added, the samples were homogenized and then centrifuged. 50 μL was applied on a TLC plate (ALUGRAM^®^ Xtra SIL G UV_254_, Macherey-Nagel). For the mobile phase CHCl_3_/MeOH in a ratio of 9:5 (v/v) with 1 % (v) formic acid was used. Visualization was performed at 254 nm (Figure 7) or spraying with anisaldehyde (Figure S3).

### Bioinformatics

Annotations of the different genes were obtained by the KEGG database (***Kanehisa and Goto, 2000***). Also, radical SAM enzyme search was done in KEGG database (searching for pf: Radical_SAM, PF04055). Searching for homologous genes was performed by using BlastP (BLOSUM 62). Searching for Ald2 homologs in KEGG database, *R. rubrum* protein sequence (rru:Rru_A0359) was used as a query sequence and an e-value <10e-20 was used for positive results.

## Abbreviations

SAM: *S*-Adenosylmethionine
MTA: Methylthioadenosine
5dAdo: 5-Deoxyadenosine
MSP: Methionine salvage pathway
5dR: 5-Deoxyribose
7dSh: 7-Deoxysedoheptulose
5dR-1P: 5-Deoxyribose 1-phosphate
5dRu-1P: 5-Deoxyribulose 1-phosphate
MTRI: Methylthioribose 1-phosphate isomerase
MTR: Methylthioribose

## Author Contributions

J. R. designed, performed, interpreted experiments, and wrote the manuscript. P. R. synthesized labelled and unlabelled 5dR and 7dSh. J. K. optimized the GC-MS method and supported with GC-MS measurements. K. B supported initial experiments and proof-read manuscript. S. G. supported chemical analytics and proof-read manuscript. K. F. supervised the study and supported manuscript writing.

## Acknowledgements

Work of the authors is supported and funded by the “Glycobiotechnology” initiative (Ministry for Science, Research and Arts Baden-Württemberg), the RTG 1708 “Molecular principles of bacterial survival strategies” and the Institutional Strategy of the University of Tübingen (Deutsche Forschungsgemeinschaft, ZUK 63). The work was further supported by infrastructural funding from the DFG Cluster of Excellence EXC 2124 Controlling Microbes to Fight Infections. We thank Dr. Libera Lo Presti for critical reading the manuscript. We especially thank Tim Orthwein for fruitful discussions and Michaela Schuppe for the cultivation of *Streptomyces setonensis*.

