## Supplementary information for "5-Deoxyadenosine Salvage by Promiscuous Enzyme Activity leads to Bioactive Deoxy-Sugar Synthesis in *Synechococcus elongatus*"

### Results

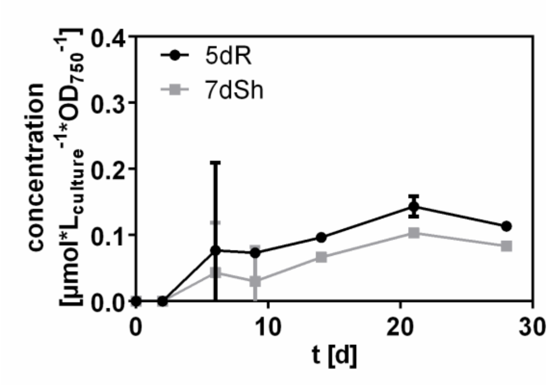


**Figure S1: Only small intracellular 5dR and 7dSh accumulation in *S. elongatus.*** Concentration of 5dR (black dots) and 7dSh (grey squares) in *S. elongatus* cells [µmol*$L^{-1}_{culture}$*$OD_{750}^{-1}$] aerated with air supplemented with 2 % $CO_2$.

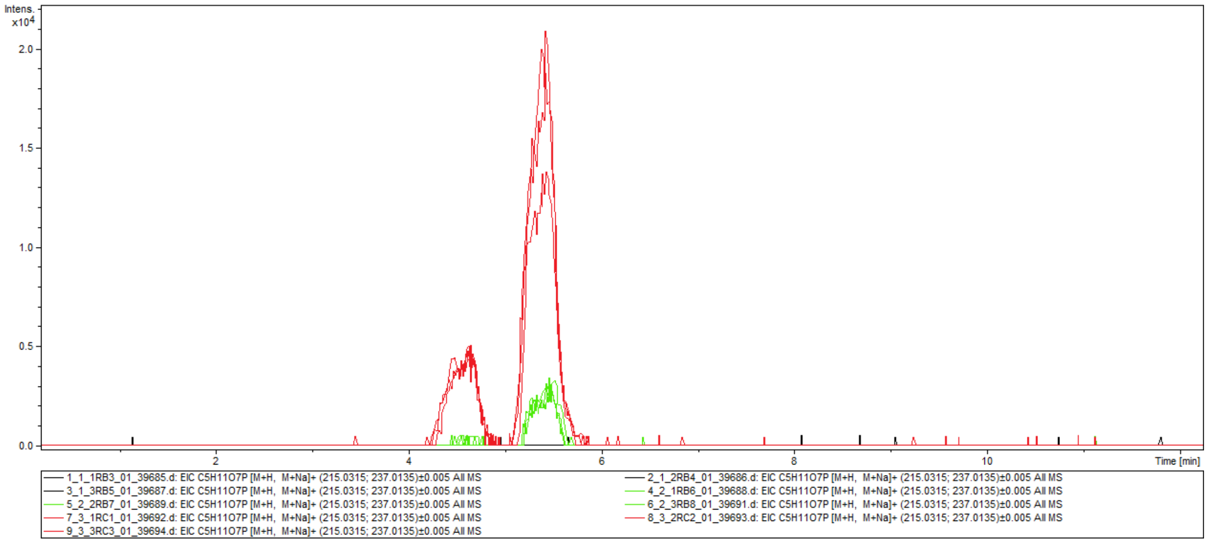


**Figure S2: 5dR-1P accumulates in crude extracts of *S. elongatus* that were incubated in the presence of phosphate.** Accumulation of 5dR-1P shown as extracted ion chromatogram [M+H, M+Na]+ (*m/z* 215.0315; *m/z* 237.0135) in crude extracts of *S. elongatus*, (Red – with 5dAdo+potassium phosphate buffer (PPB); green – with 5dAdo, no PPB; black – without 5dAdo, PPB). Three independent replicates are shown for each treatment. One part of the samples of the crude extract assays was analysed via high resolution LC-MS (C18 Gemini, solvent A: ACN+0.1 %TFA, solvent B: $H_2O$, 1% - 20% B in 20 min, Maxis 4G ESI-QTOF).

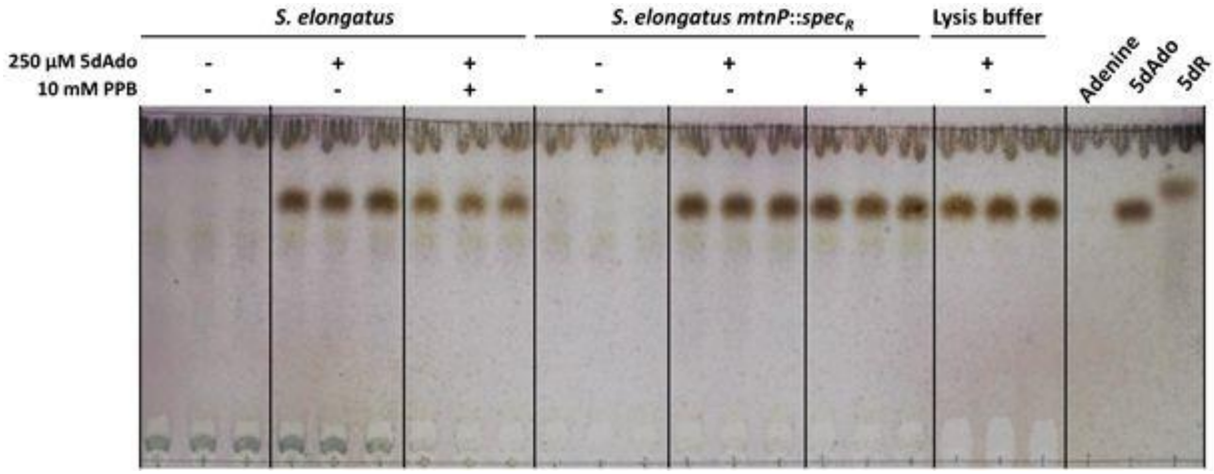


**Figure S3: 5dR does not accumulate in crude extracts which were incubated with 5dAdo.** Crude extracts from *S. elongatus* or *S. elongatus mtnP*::*spec$_R$* were incubated with 5dAdo in the presence or absence of potassium phosphate buffer (PPB) and then analysed via thin layer chromatography (TLC). TLC plate from Figure 7 (main text) was sprayed with anisaldehyde after UV-visualisation. Pure adenine, 5dAdo and 5dR were used as standards (right). Adenine is only visible with UV-visualisation (see Figure 7, main text). Three independent replicates are shown for each condition.

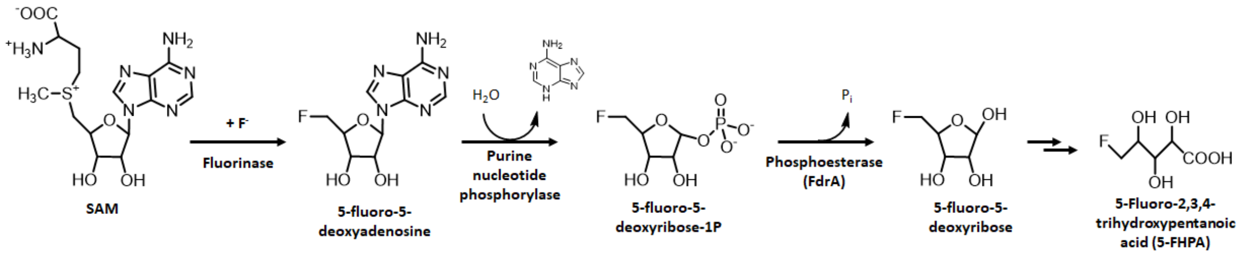


**Figure S4: Biosynthesis of the fluoro-metabolite 5-FHPA in *Streptomyces* sp. MA37 (modified after *Ma et al.* (*2015*)).**

**Table S1: Overview of MSP genes present in different cyanobacteria.**

| Gene name in *B. subtilis* | | MtnN | MtnK | MtnP* | MtnA | MtnB | MtnC*** | MtnX | MtnW | MtnD | MtnE |
|---|---|---|---|---|---|---|---|---|---|---|---|
| **Gene Product** | | **MTA nucleosidase** | **MTR kinase** | **MTA phosphory-lase** | **MTR-1P isomerase** | **MTRu-1P dehydratase **** | **enolase/ phosphatase[*1]** | **phospha-tase[*2]** | **enolase[*3]** | **dioxygenase[*4]** | **amino transferase ****** |
| Strain | Identifier | EC 3.2.2.9 | EC 2.7.1.100 | EC 2.4.2.28 | EC 5.3.1.23 | EC 4.2.1.109 | EC 3.1.3.77 | EC 3.1.3.87 | EC 5.3.2.5 | EC 1.13.11.53/54 | EC 2.6.1.117 |
| PCC 7942 | syf: Synpcc7942_ | - | - | 0923 | 1992 | 1993 | 1994 | 0510 | - | 0608 | 2.6.1.- |
| PCC 6301 | syc: syc | - | - | 0619_d | 2104_d | 2103_c | 2102_c | 1010_d | - | 0916_c | 2.6.1.- |
| PCC 7002 | syp: SYNPCC7002_ | - | - | A0108 | A2308 | A0554 | A0552 | A0257 | - | A0553 | 2.6.1.- |
| PCC 6803 | syn: | - | - | sll0135 | slr1938 | - | - | - | - | - | 2.6.1.- |
| PCC 7120 | ana: | - | - | alr4054 | all3566 | - | - | - | - | all2724 | 2.6.1.- |
| PCC 7502 | synp: Syn7502_ | - | - | 03055 | 00983 | - | - | - | - | - | 2.6.1.- |
| PCC 6312 | syne: Syn6312_ | - | - | 2991 | 2219 | - | - | - | - | - | 2.6.1.- |
| ATCC 29431 | ava: | - | - | Ava_1653 | Ava_3544 | - | - | - | - | Ava_4291 | 2.6.1.- |
| ATCC 29133 | npu: Npun_ | - | - | F6610 | F5471 | - | F4952 | - | - | - | 2.6.1.- |

Gene abbreviations according to *B. subtilis* annotation in which the MSP was discovered (***Sekowska et al., 2004; Sekowska and Danchin, 2002***). Gene identifiers are referred to KEGG (***Kanehisa and Goto, 2000***).

* not in *B. subtilis*

** For *A. thaliana* MTR-1P dehydratase (DEP1) moonlighting aldolase activity on 5-deoxyribulose 1-phosphate was shown (***Beaudoin et al., 2018,*** Supporting information).

*** not in *B. subtilis*. MtnC (EC 3.1.3.77) is a bifunctional enzyme, which has enolase and phosphatase activity. In *B. subtilis* this step is performed by two enzymes (MtnW: 2,3-diketo-5-methylthiopentyl-1-phosphate enolase (EC 5.3.2.5) and MtnX: 2-hydroxy-3-keto-5-methylthiopentenyl-1-phosphate phosphatase (EC 3.1.3.87).

**** aminotransferases are normally broad specificity enzymes (***Sekowska et al., 2004***)

[*1] 2,3-dioxomethio-pentane-1P enolase/phosphatase

[*2] 2-hydroxy-3-keto-5-methylthiopentenyl-1-phosphate phosphatase

[*3] 2,3-diketo-5-methylthiopentyl-1-phosphate enolase

[*4] 1,2-dihydroxy-3-keto-5-methylthiopentene dioxygenase

**Table S2: Genes encoding for SAM radical enzymes in *S. elongatus* PCC 7942.** KEGG genes were examined for the presence of the Pfam motif PF04055 (SAM_radical), which is a distinctive feature of radical SAM enzymes. Table shows accession number and annotations from KEGG. GenBank annotations are only shown if using another annotation.

| Accesion No. | Annotation |
|---|---|
| Synpcc7942_0419 | K01012 biotin synthase [EC:2.8.1.6] |
| Synpcc7942_0542 | K03644 lipoyl synthase [EC:2.8.1.8] |
| Synpcc7942_0686 | K11781 5-amino-6-(D-ribitylamino)uracil---L-tyrosine 4-hydroxyphenyl transferase [EC:2.5.1.147] \| (GenBank) FO synthase subunit 2 |
| Synpcc7942_0799 | no KO assigned \| (GenBank) Elongator protein 3 |
| Synpcc7942_0838 | no KO assigned \| (GenBank) Elongator protein 3/MiaB/NifB |
| Synpcc7942_0877 | no KO assigned \| (GenBank) Elongator protein 3/MiaB/NifB |
| Synpcc7942_0892 | K11780 7,8-didemethyl-8-hydroxy-5-deazariboflavin synthase [EC:4.3.1.32] \| (GenBank) FO synthase subunit 1 |
| Synpcc7942_0945 | no KO assigned \| (GenBank) conserved hypothetical protein |
| Synpcc7942_1229 | K05936 precorrin-4/cobalt-precorrin-4 C11-methyltransferase [EC:2.1.1.133 2.1.1.271] |
| Synpcc7942_1282 | K03639 GTP 3',8-cyclase [EC:4.1.99.22] \| (GenBank) GTP cyclohydrolase subunit MoaA |
| Synpcc7942_1332 | K10026 7-carboxy-7-deazaguanine synthase [EC:4.3.99.3] \| (GenBank) conserved hypothetical protein |
| Synpcc7942_1507 | K03644 lipoyl synthase [EC:2.8.1.8] \| (GenBank) lipoic acid synthetase |
| Synpcc7942_1621 | no KO assigned \| (GenBank) Elongator protein 3/MiaB/NifB |
| Synpcc7942_1652 | no KO assigned \| (GenBank) Elongator protein 3/MiaB/NifB |
| Synpcc7942_1758 | K06941 23S rRNA (adenine2503-C2)-methyltransferase [EC:2.1.1.192] \| (GenBank) conserved hypothetical protein |
| Synpcc7942_2374 | K06168 tRNA-2-methylthio-N6-dimethylallyladenosine synthase [EC:2.8.4.3] \| (GenBank) tRNA-i(6)A37 thiotransferase enzyme MiaB |
| Synpcc7942_2382 | no KO assigned \| (GenBank) coproporphyrinogen III oxidase, anaerobic |
| Synpcc7942_2512 | K14441 ribosomal protein S12 methylthiotransferase [EC:2.8.4.4] \| (GenBank) Protein of unknown function UPF0004 |

**Table S3: Genes encoding for phosphoric monoester hydrolases [EC: 3.1.3.-] in *Synechococcus elongatus* PCC 7942.** Table shows accession number and annotations from KEGG. GenBank annotations are only shown if using another annotation.

| Accesion No. | Annotation |
|---|---|
| Synpcc7942_0173 | K01082 3'(2'), 5'-bisphosphate nucleotidase [EC:3.1.3.7] \| (GenBank) 3'-Phosphoadenosine 5'-phosphosulfate (PAPS) 3'-phosphatase-like |
| Synpcc7942_0463 | K01104 protein-tyrosine phosphatase [EC:3.1.3.48] \| (GenBank) protein tyrosine phosphatase |
| Synpcc7942_0485 | K22305 phosphoserine phosphatase [EC:3.1.3.3] \| (GenBank) phosphoglycerate mutase |
| Synpcc7942_0505 | K11532 fructose-1,6-bisphosphatase II / sedoheptulose-1,7-bisphosphatase [EC:3.1.3.11 3.1.3.37] |
| Synpcc7942_0510 | K08966 2-hydroxy-3-keto-5-methylthiopentenyl-1-phosphate phosphatase [EC:3.1.3.87] |
| Synpcc7942_0613 | K08296 phosphohistidine phosphatase [EC:3.1.3.-] \| (GenBank) phosphohistidine phosphatase, SixA |
| Synpcc7942_0693 | K01091 phosphoglycolate phosphatase [EC:3.1.3.18] \| (GenBank) conserved hypothetical protein |
| Synpcc7942_0791 | K00974 tRNA nucleotidyltransferase (CCA-adding enzyme) [EC:2.7.7.72 3.1.3.- 3.1.4.-] \| (GenBank) polyA polymerase |
| Synpcc7942_0965 | K01082 3'(2'), 5'-bisphosphate nucleotidase [EC:3.1.3.7] \| (GenBank) ammonium transporter protein Amt1-like |
| Synpcc7942_0976 | K00974 tRNA nucleotidyltransferase (CCA-adding enzyme) [EC:2.7.7.72 3.1.3.- 3.1.4.-] \| (GenBank) CBS |
| Synpcc7942_1005 | K20866 glucose-1-phosphatase [EC:3.1.3.10] \| (GenBank) HAD-superfamily hydrolase subfamily IA, variant 3 |
| Synpcc7942_1130 | K01090 protein phosphatase [EC:3.1.3.16] \| (GenBank) protein serine/threonine phosphatase |
| Synpcc7942_1515 | K01090 protein phosphatase [EC:3.1.3.16] \| (GenBank) protein serine/threonine phosphatase |
| Synpcc7942_1553 | K07053 3',5'-nucleoside bisphosphate phosphatase [EC:3.1.3.97] \| (GenBank) Phosphoesterase PHP-like |
| Synpcc7942_1763 | K01092 myo-inositol-1(or 4)-monophosphatase [EC:3.1.3.25] \| (GenBank) inositol monophosphate family protein |
| Synpcc7942_1931 | K07313 serine/threonine protein phosphatase 1 [EC:3.1.3.16] \| (GenBank) probable serine/threonine protein phosphatase |
| Synpcc7942_1994 | K09880 enolase-phosphatase E1 [EC:3.1.3.77] \| (GenBank) 2,3-diketo-5-methylthio-1-phosphopentane phosphatase |
| Synpcc7942_2063 | K03787 5'-nucleotidase [EC:3.1.3.5] \| (GenBank) exopolyphosphatase / 5'-nucleotidase / 3'-nucleotidase |
| Synpcc7942_2076 | K06949 ribosome biogenesis GTPase / thiamine phosphate phosphatase [EC:3.6.1.- 3.1.3.100] \| (GenBank) GTPase EngC |
| Synpcc7942_2288 | K03270 3-deoxy-D-manno-octulosonate 8-phosphate phosphatase (KDO 8-P phosphatase) [EC:3.1.3.45] \| (GenBank) Phosphatase kdsC |
| Synpcc7942_2335 | K03841 fructose-1,6-bisphosphatase I [EC:3.1.3.11] \| (GenBank) D-fructose 1,6-bisphosphatase |
| Synpcc7942_2473 | K07315 phosphoserine phosphatase RsbU/P [EC:3.1.3.3] \| (GenBank) serine phosphatase |
| Synpcc7942_2582 | K01092 myo-inositol-1(or 4)-monophosphatase [EC:3.1.3.25] |
| Synpcc7942_2589 | K05979 2-phosphosulfolactate phosphatase [EC:3.1.3.71] |
| Synpcc7942_2613 | K01091 phosphoglycolate phosphatase [EC:3.1.3.18] \| (GenBank) HAD-superfamily hydrolase subfamily IA |

### Material & Methods

**Table S4: Oligonucleotides for PCR amplification (G) and sequencing (S). Overlapping fragments for Gibson cloning are labelled in red.**

| Name | Sequence (5‘ → 3‘) |
|---|---|
| 46_0923_up_fw (G) | AGCTCGGTACCCGGGGATCCTGGCCTCTACCGAATGGAAGC |
| 47_0923_up_rev (G) | CTGCGTTCGGTCAAGAGCTTTGCCAAAGAAGGTCGAAGG |
| 32_Spec_fw (G) | GAGCTCTTGACCGAACGCAG |
| 33_Spec_rev (G) | TTATTTGCCGACTACCTTGGTGATCTC |
| 48_0923_down_fw (G) | GAGATCACCAAGGTAGTCGGCAAATAACCCAGATTATCGGCATGACC |
| 49_0923_down_rev (G) | ACGCCAAGCTTGCATGCCTGCATGCAGGAGTAGTGCCAAACG |
| 1064_pUC19_fw (S) | TGCTGCAAGGCGATTAAGTTGGG |
| 1065_pUC19_rev (S) | CGACAGGTTTCCCGACTGGAAAG |
| 50_0923_rev_seg (S) | CTAGTCGACCCGCTTCAACC |
| 51_0923_fw_seg (S) | GCCACCAAGGATCCCAGATG |
| 85_1005_up_fw | AGCTCGGTACCCGGGGATCCTTACAACCGCCTCAAGTGC |
| 86_1005_up_rev | CTGCGTTCGGTCAAGAGCTATGGAGCGTCCCGAAGTAAG |
| 87_1005_down_fw | GAGATCACCAAGGTAGTCGGCAAATAAATGCTTGCTCGTCTTGG |
| 88_1005_down_rev | ACGCCAAGCTTGCATGCCTGCAGCTGCTCCAAAGGCAAAC |

#### Chemical synthesis of 5-deoxyribose and 7-deoxysedoheptulose

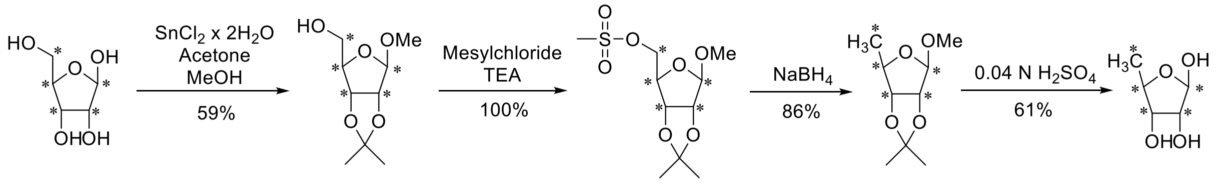


(1) (2) (3) (4) (5)

**Figure S5: Four step synthesis of [U-$^{13}C_5$]-5dR modified after(*Sairam et al., 2003; Zhang et al., 2013*). (1): $^{13}C_5$-D-Ribose, (2): Methyl-2,3-*O*-isopropylidene-$^{13}C_5$-*β*-D-ribofuranoside, (3): Methyl-2,3-*O*-isopropylidene-5-*O*-mesyl-$^{13}C_5$-*β*-D-ribofuranoide, (4): Methyl-2,3-*O*-isopropylidene-$^{13}C_5$-5-deoxy-*β*-D-ribofuranoide, (5): [U-$^{13}C_5$]-5-Deoxy-D-ribofuranose.**

##### Physiochemical data of intermediates of the chemical synthesis

**Abbreviations:** TLC: thin layer chromatography; $R_f$: retention factor; NMR: nuclear magnetic resonance; MHz: megahertz; $CDCl_3$: deuterated chloroform; δ: chemical shift; ppm: part per million; dm: doublet of multiplet; *J*: coupling constants; Hz: hertz; ddm: doublet of doublet of multiplet; d: doublet; s: singlet; m: multiplet; HR-(+)ESI-MS: High resolution-electrospray-mass spectrometry (positive mode); m/z: mass-to-charge ratio; calcd: calculated; ddd: doublet of doublet of doublet; $D_2O$; deuterium oxide.

##### Methyl-2,3-*O*-isopropylidene-$^{13}C_5$-β-D-ribofuranoside (2)

**TLC:** $R_f$ 0.63 (cyclohexane/ethylacetate 1:1)

**$^1$H-NMR** (400 MHz, $CDCl_3$):
δ (ppm)=4.95 (dm, $J_{1,C\text{-}1}$=174.4 Hz, 1H, 1-H), 4.81 (dm, $J_{3,C\text{-}3}$=160.6 Hz, 1H, 3-H), 4.57 (dm, $J_{2,C\text{-}2}$=158.7 Hz, 1H, 2-H), 4.41 (dm, $J_{4,C\text{-}4}$=155.1 Hz, 1H, 4-H), 3.68 (ddm, $J_{5a,C\text{-}5}$=143.8 Hz, $J_{5a,5b}$=12.6 Hz, 1H, 5a-H), 3.59 (ddm, $J_{5b,C\text{-}5}$=142.0 Hz, $J_{5b,5a}$=12.6 Hz, 1H, 5b-H), 3.42 (d, $J_{\text{-}OCH3,C\text{-}1}$=4.4 Hz, 3H, -O*CH₃*), 1.47 and 1.30 (s, 3H, C*(CH₃)₂*)

**$^{13}$C-NMR** (100 MHz, $CDCl_3$):
δ (ppm)=112.2 (*C*($CH_3$)$_2$), 109.8 (d, $J_{C\text{-}1,C\text{-}2}$=48.5 Hz, C-1), 88.4 (dd, $J_{C\text{-}4,C\text{-}3}$=39.8 Hz, $J_{C\text{-}4,C\text{-}5}$=38.9 Hz, C-4), 85.9 (dd, $J_{C\text{-}2,C\text{-}1}$=48.5 Hz, $J_{C\text{-}2,C\text{-}3}$=31.0 Hz, C-2), 81.5 (ddm, $J_{C\text{-}3,C\text{-}4}$=39.8 Hz, $J_{C\text{-}3,C\text{-}2}$=31.0 Hz, C-3), 64.1 (dm, $J_{C\text{-}5,C\text{-}4}$=38.9 Hz, C-5), 55.6 (m, -O*CH₃*), 26.4 and 24.8 (C*(CH₃)₂*)

**HR-(+)ESI-MS:** *m/z* calcd. for $[M+H]^+$: 210.1238, found: 210.1239; *m/z* calcd. for $[M+Na]^+$: 232.1058, found: 232.1058.

##### Methyl-2,3-*O*-isopropylidene-5-*O*-mesyl-$^{13}C_5$-β-D-ribofuranoide (3)

**TLC:** $R_f$ 0.59 (cyclohexane/ethylacetate 1:1)

**$^1$H-NMR** (400 MHz, $CDCl_3$):
δ (ppm)=4.98 (ddd, $J_{1,C\text{-}1}$=173.9 Hz, $J_{1,C\text{-}2}$=7.5 Hz, $J_{1,2}$=3.0 Hz, 1H, 1-H), 4.69 (dm, $J_{3,C\text{-}3}$=154.5 Hz, 1H, 3-H), 4.60 (ddm, $J_{2,C\text{-}2}$=158.9 Hz, $J_{2,1}$=3.0 Hz, 1H, 2-H), 4.40 (dm, $J_{4,C\text{-}4}$=155.1 Hz, 1H, 4-H), 4.20 (dm, $J_{5a,C\text{-}5}$=152.1 Hz, 1H, 5a-H), 4.18 (dm, $J_{5b,C\text{-}5}$=154.1 Hz, 1H, 5b-H), 3.34 (d, $J_{\text{-}OCH3,C\text{-}1}$=4.5 Hz, 3H, -O*CH₃*), 3.06 (s, 1H, Mesyl-$CH_3$), 1.47 and 1.30 (s, 3H, C*(CH₃)₂*)

**$^{13}$C-NMR** (100 MHz, $CDCl_3$):

δ (ppm)=113.0 (*C*($CH_3$)$_2$), 109.7 (d, $J_{C-1,C-2}$=49.3 Hz, C-1), 85.0 (dd, $J_{C-2,C-1}$=49.3 Hz, $J_{C-2,C-3}$=30.7 Hz, C-2), 83.9 (dd, $J_{C-4,C-5}$=42.5 Hz, $J_{C-4,C-3}$=39.1 Hz, C-4), 81.5 (ddd, $J_{C-3,C-4}$=39.1 Hz, $J_{C-3,C-2}$=30.7 Hz, $J_{C-3,C-5}$=5.5 Hz, C-3), 68.5 (dd, $J_{C-5,C-4}$=42.5 Hz, $J_{C-5,C-3}$=5.5 Hz, C-5), 55.4 (m, -O$CH_3$), 37.9 (Mesyl-$CH_3$), 26.5 and 25.0 (C$(CH_3)_2$)

**HR-(+)ESI-MS:** *m/z* calcd. for $[M+H]^+$: 288.1014, found: 288.1013; *m/z* calcd. for $[M+Na]^+$: 310.0833, found: 310.0831.

##### Methyl-2,3-*O*-isopropylidene-$^{13}C_5$-5-deoxy-β-D-ribofuranoide (4)

**TLC:** $R_f$ 0.87 (cyclohexane/ethylacetate 1:1)

**$^1$H-NMR** (700 MHz, $CDCl_3$):
δ (ppm)=4.92 (ddd, $J_{1,C-1}$=172.3 Hz, $J_{1,C-2}$=7.5 Hz, $J_{1,2}$=2.6 Hz, 1H, 1-H), 4.61 (ddm, $J_{2,C-2}$=161.4 Hz, $J_{2,1}$=2.6 Hz, 1H, 2-H), 4.49 (dm, $J_{3,C-3}$=155.8 Hz, 1H, 3-H), 4.32 (dm, $J_{4,C-4}$=149.6 Hz, 1H, 4-H), 3.31 (d, $J_{-OCH3,C-1}$=4.4 Hz, 3H, -O$CH_3$), 1.46 and 1.29 (s, 3H, C$(CH_3)_2$), 1.27 (dm, $J_{5,C-5}$=126.3 Hz, 3H, 5-H)

**$^{13}$C-NMR** (176 MHz, $CDCl_3$):
δ (ppm)=112.2 (*C*($CH_3$)$_2$), 109.6 (dm, $J_{C-1,C-2}$=48.1 Hz, C-1), 85.9 (dm, $J_{C-2,C-1}$=48.1 Hz, C-2), 85.3 (dm, $J_{C-3,C-4}$=37.7 Hz, C-3), 83.2 (ddm, $J_{C-4,C-3}$=$J_{C-4,C-5}$=37.7 Hz, C-4), 54,5 (m, -O$CH_3$), 26.6 and 25.1 (C$(CH_3)_2$), 21.0 (dm, $J_{C-5,C-4}$=37.7 Hz, C-5)

**HR-(+)ESI-MS:** *m/z* calcd. for $[M+H]^+$: 194.1289, found: 194.1294; *m/z* calcd. for $[M+Na]^+$: 216.1109, found: 216.1111.

##### [U-$^{13}C_5$]-5-Deoxy-D-ribofuranose (5)

**TLC:** $R_f$ 0.46 (chloroform/methanol 4:1)

**$^1$H-NMR** (400 MHz, $D_2O$):
β-furanose: δ (ppm)=5.18 (dm, $J_{1,C-1}$=172.1 Hz, 1H, 1-H), 4.00-3.95 (m, 3H, 2-H, 3-H, 4-H), 1.33 (dm, $J_{3,C-3}$=126.8 Hz, 3H, 5-H)
α-furanose: δ (ppm)= 5.35 (dm, $J_{1,C-1}$=172.6 Hz, 1H, 1-H), 4.14 (dm, $J_{2,C-2}$=151.2 Hz, 1H, 2-H), 4.12 (dm, $J_{4,C-4}$=150.5 Hz, 1H, 4-H), 3.80 (dm, $J_{3,C-3}$=150.2 Hz 1H, 3-H), 1.24 (dm, $J_{3,C-3}$=126.8 Hz, 3H, 5-H)

**$^{13}$C-NMR** (100 MHz, $D_2O$):
β-furanose: δ (ppm)= 100.8 (m, C-1), 78.3 (dm, $J_{C-4,C-5}$=39.5 Hz, C-4), 75.2 (m, C-2 and C-3), 19.1 (d, $J_{C-5,C-4}$=39.5 Hz, C-5)
α-furanose: δ (ppm)= 95.7 (dm, $J_{C-1,C-2}$=42.2 Hz, C-1), 78.1 (dm, $J_{C-4,C-5}$=39.9 Hz, C-4), 74.8 (m, C-3), 70.4 (dm, $J_{C-2,C-1}$=42.2 Hz, C-2), 17.8 (d, $J_{C-5,C-4}$=39.9 Hz, C-5)

**HR-(+)ESI-MS:** *m/z* calcd. for $[M+Na]^+$: 162.0639, found: 162.0640.

##### [3,4,5,6,7-$^{13}C_5$]-7-Deoxy-D-*altro*-heptulose

**TLC:** $R_f$ 0.56 (chloroform/methanol 8:5)

**$^1$H-NMR** (400 MHz, $D_2O$):
β-furanose δ (ppm)=4.21 (dm, $J_{4,C-4}$=110.7 Hz, 1H, 4-H), 4.07 (dm, $J_{3,C-3}$=145.4 Hz, 1H, 3-H), 3.94 (m, 1H, H-6), 3.69 (m, 1H, 5-H), 3.63 (dd, $J_{1a,1b}$=11.8 Hz, $J_{1a,C-3}$=4.3 Hz, 1H, 1a-H), 3.54 (d, $J_{1b,1a}$=11.8 Hz, $J_{1b,C-3}$=6.5 Hz, 1H, 1b-H), 1.20 (dm, $J_{7,C-7}$=126.8 Hz, 3H, 7-H)
α-pyranose δ (ppm)=4.07 (m, 1H, 6-H), 4.03 (m, 1H, 4-H), 3.69 (m, 1H, 3-H), 3.66 (m, 1H, 1a-H), 3.57 (m, 1H, 5-H), 3.40 (m, 1H, 1b-H), 1.26 (dm, $J_{7,C-7}$=127.1 Hz, 3H, 7-H)
α-furanose δ (ppm)=4.14 (m, 1H, 4-H), 4.06 (m, 1H, 3-H), 3.99 (m, 1H, 6-H), 3.91 (m, 1H, 5-H), 3.92 (m, 1H, 1a-H), 3.64 (m, 1H, 1b-H), 1.20 (dm, $J_{7,C-7}$=126.8 Hz, 3H, 7-H)

**$^{13}$C-NMR** (100 MHz, $D_2O$):

β-furanose δ (ppm)=101.3 (dm, $J_{C\text{-}2,C\text{-}3}$=44.4 Hz, C-2), 83.5 (dm, $^{1}J$=43.4 Hz, 39.0 Hz, $^{2}J$=5.4 Hz, C-5), 75.8 (m, C-3); 74.7 (m, C-4), 67.7 (m, C-6), 62.4 (C-1), 17.0 (d, $J_{C\text{-}7,C\text{-}6}$=38.4 Hz, C-7)

α-pyranose δ (ppm)=98.0 (m, C-2), 70.8 (dm, $^{1}J$=38.9 Hz, C-4), 68.9 (m, C-5), 67.6 (m, C-3), 64.5 (dm, $^{1}J$=41.1 Hz, C-6), 63.8 (C-1), 16.9 (m, C-7)

α-furanose δ (ppm)= 104.4 (m, C-2), 84.7 (dm, $^{1}J$=40.9 Hz, C-5), 82.0 (dm, $J_{C\text{-}3,C\text{-}4}$=40.5 Hz, C-3), 75.8 (m, C-4), 66.7 (dm, $^{1}J$=38.4 Hz, C-6), 62.9 (m, C-1), 17.0 (d, $J_{C\text{-}7,C\text{-}6}$=37.4 Hz, C-7)

**HR-(+)ESI-MS:** *m/z* calcd. for $[M+Na]^{+}$: 222.0850, found: 222.0852.
